# Dynamic chromosome rearrangements of the white-spotted bamboo shark shed light on cartilaginous fish diversification mechanisms

**DOI:** 10.1101/602136

**Authors:** Yaolei Zhang, Haoyang Gao, Hanbo Li, Jiao Guo, Meiniang Wang, Qiwu Xu, Jiahao Wang, Meiqi Lv, Xinyu Guo, Qun Liu, Likun Wei, Han Ren, Yang Xi, Yang Guo, Qian Zhao, Shanshan Pan, Chuxin Liu, Liping Sang, Xiaoyan Ding, Chen Wang, Haitao Xiang, Yue Song, Yujie Liu, Shanshan Liu, Yuan Jiang, Changwei Shao, Jiahai Shi, Shiping Liu, Jamal S. M. Sabir, Mumdooh J. Sabir, Muhummadh Khan, Nahid H. Hajrah, Simon Ming-Yuen Lee, Xun Xu, Huanming Yang, Jian Wang, Guangyi Fan, Naibo Yang, Xin Liu

## Abstract

Cartilaginous fishes have a very high phenotypical diversity, a phenomenon for which the mechanisms have been largely unexplored. Here, we report the genome of the white-spotted bamboo shark as the first chromosome-level genome assembly of cartilaginous fish. Using this genome, we illustrated a dynamic chromosome rearrangement process in the bamboo shark, which resulted in the formation of 13 chromosomes, all of which were sparsely distributed with conserved genes and fast-evolving. We found the fast-evolving chromosomes to be enriched in immune-related genes with two chromosomes harboring the major genes for developing the single-chain antibody. We also found chromosome rearrangements to have resulted in the loss of two genes (*p2rx3* and *p2rx5*) which we also showed were involved in cartilage development using a CRISPR/Cas9 approach in zebrafish. Our study highlighted the significance of chromosome rearrangements in the phenotypical evolution of cartilaginous fishes, providing clues to inform further studies on mechanisms for fish diversification.

## Introduction

The white-spotted bamboo shark, *Chiloscyllium plagiosum*, (hereinafter referred to as bamboo shark) belongs to the class of Chondrichthyes, which is one of the oldest extant jawed vertebrate groups ^1^. The cartilaginous fishes diverged from a common vertebrate ancestor about 460-520 million years ago (MYA), and have been since evolving independently into distinct lineages ^2^. Most cartilaginous fishes have a large number of chromosomes and this number varies between species (2n= 66∼104) ^3,4^, but the chromosomal evolution of cartilaginous fishes has remained largely unexplored. To better understand the evolution of these fishes, we sequenced and assembled the genome of a female bamboo shark. We then constructed a chromosome level reference genome, identified dynamic chromosome rearrangement events and analyzed their evolutionary consequences. Our results are innovative and significant, which are different from previous cartilaginous fish genome^5^.

## Results

We assembled a 3.85 Gb genome assembly with 51 chromosomes using a whole genome shotgun strategy combined with Hi-C sequencing data. To our knowledge, this is the first cartilaginous fish genome to be assembled at the chromosome level (with ∼95.8% of the Benchmarking Universal Single-Copy Orthologs, BUSCOs^6^, to be complete in the genome, and **Supplementary Figure 1-4 and Supplementary Table 1-6**). Syntenic relationships revealed unambiguous alignment of 41 bamboo shark chromosomes to 29 chicken chromosomes (**Supplementary Figure 5**), while alignment between bamboo shark chromosomes and zebrafish chromosomes indicated a number of intricate matchups (**Supplementary Figure 6**), an observation which was consistent with the previous finding of extensive interchromosomal rearrangements in zebrafish^7^.

We then reconstructed the ancestral chromosomes of cartilaginous fishes by identifying and inspecting the paralogous and orthologous genes between the bamboo shark genome and the elephant shark genome^8^ following a previously described method^9^ (**Supplementary Table 7**). Finally, we constructed 21 putative ancestral chromosomes and illustrated an evolutionary scenario in which eight fission and five fusion events occurred (color arrows), possibly for all sharks (**Fig. 1a**). As for the bamboo shark, nine fission and four fusion events (black and dotted arrows) occurred, resulting in six candidate daughter chromosomes (Chr8, Chr29, Chr38 and Chr39, Chr45, Chr48) (**Fig. 1a, Supplementary Table 7** and **Supplementary Figure 7**). All of these rearrangements ultimately resulted in the 51 chromosomes of the bamboo shark genome along with individual gene gains and losses.

**Fig. 1.**
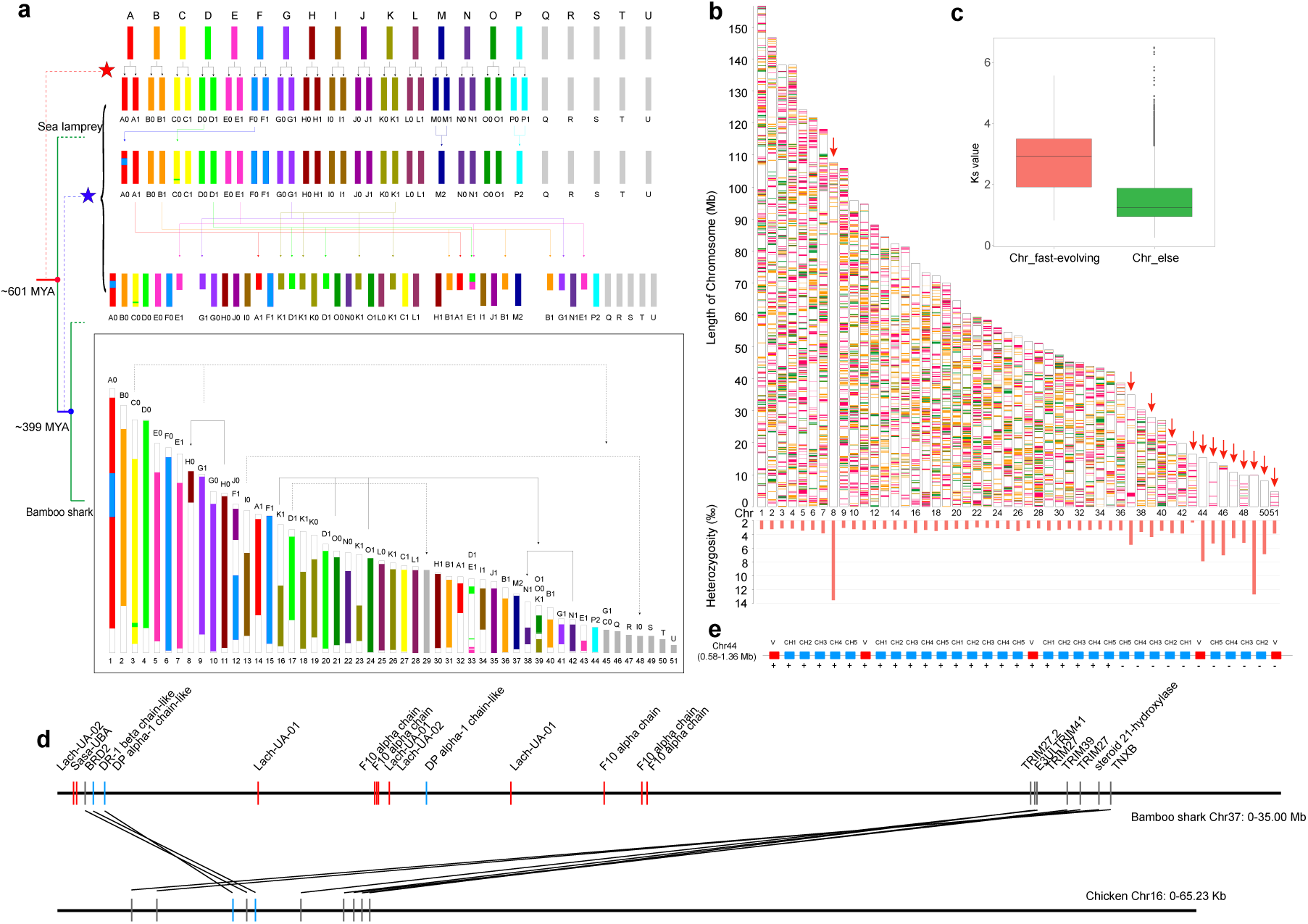
Chromosome evolution of bamboo shark. **a)** Construction of the ancestral chromosome model of elephant shark and bamboo shark. Letters from A to U represent constructed ancestral chromosomes. The grey bars represent chromosomes which did not experience duplication in the present study. The colored arrows represent rearrangements of chromosomes. The black arrows represent rearrangements specific to the bamboo shark. The dotted arrows represent potential rearrangements which are supported by gene pairs. The red star represents whole genome duplication occurred before the divergence of sea lamprey and gnathostome^52^ (early than ∼601 million years ago). The blue star represents rearrangements of shark ancestor. The times were referred to TimeTree database. **b)** Distribution of conserved genes of elephant shark, bamboo shark, whale shark and medaka. Magenta, conserved regions shared by four species (R1). Orange, common regions in three shark genome excluding R1. Green, regions shared between the white-spotted bamboo shark and medaka excluding R1. The red arrows point out 13 fast-evolving chromosomes. **c)** up: Comparison of *Ks* values of single copy orthologous genes between 13 fast-evolving chromosomes and other chromosomes. down: Heterozygosity of 51 chromosomes in bamboo shark genome. **d)** Distribution of MHC genes on chromosome 37. The red and blue rectangles represent MHC class I and class II genes, respectively. The grey rectangles represent non-MHC genes. Here, we only show syntenic genes and shark MHC genes compared with chicken. **e)** Distribution of identified IgNAR loci on chromosome 44.

To identify the potential causes and consequences of dynamic chromosome rearrangements in cartilaginous fishes, we further analyzed the distribution of conserved genes in cartilaginous fishes along the bamboo shark chromosomes. We identified 6,205 orthologous genes (∼31.66% of total genes) shared by the bamboo shark, elephant shark and whale shark (**Supplementary Figure 8**). After exclusion of the genes shared among these three cartilaginous fishes and representative bony fishes (medaka^7^, **Supplementary Figure 9** and spotted gar^10^, **Supplementary Figure 10**), we identified 3,678 genes conserved in cartilaginous fishes (**Fig. 1b**). Interestingly, we found those genes to be unevenly distributed along bamboo shark chromosomes with conserved genes on chromosomes 8, 37, 39, 41, 43, 44, 45, 46, 47, 48, 49, 50 and 51 - notably fewer than those of other chromosomes (Mann-Whitney U test, *p*-value<0.001, **Fig. 1b** and **Supplementary Table 8**). We then evaluated the evolutionary rate of genes on these chromosomes by calculating the *KS* (synonymous substitutions per synonymous site) values of 160 single copy orthologous genes on these 13 chromosomes (mean *KS* value: 2.79), which was notably higher than that of other chromosomes (mean *KS* value: 1.54) (Mann-Whitney U test, *p*-value<0.001, **Fig. 1c**). In addition, we found heterozygous SNPs in the genome of the individual we sequenced to be notably more frequent on these 13 chromosomes than other chromosomes (Mann-Whitney U test, *p*-value<0.001, **Fig. 1b**). All of these findings above suggest that these 13 chromosomes are fast-evolving. Enrichment analysis (according to the Kyoto Encyclopedia of Genes and Genomes (KEGG)-assigned gene functions and pathways) showed that genes on these 13 fast-evolving chromosomes are significantly enriched in immune-related pathways with 171 immune-related genes (Mann-Whitney U test, *p*-value<0.01, **Supplementary Table 9-10**). These include allograft rejection, antigen processing and presentation, as well as intestinal immune network for IgA production.

Among the fast-evolving chromosomes that we identified, Chr37 and Chr44 likely underwent a special self-fusion event after a possible whole-genome duplication event (**Fig. 1a**). We found that major histocompatibility complex (MHC) genes (11 class I and 3 class II genes) are notably enriched on Chr37 (**Fig. 1d**), only except those on unanchored scaffold sequences. MHC genes were not found in the amphioxus genome while one fragment of a possible MHC class II gene was found in sea lamprey^11^. Upon further investigation of MHC gene numbers in other species, we found both MHC class I and class II genes in cartilaginous fishes and bony fishes except for the elephant shark, which lacked MHC class II genes according to our analysis (**Supplementary Table 11-13** and **Supplementary Figure 11**). These results suggest that the innate immune system played a major role in defending against infections in amphioxus and sea lamprey, while cartilaginous and bony fishes evolved to acquire a complete MHC-based, adaptive immune system. The differences in these immune systems may have arisen from the fast-evolving chromosomes. Moreover, we suggest that MHC class II genes were likely acquired before MHC class I genes based on our identification of an MHC class II-like fragment in the starlet sea anemone and sea lamprey^11^, potentially resolving a long debate about MHC evolution^12–18^. In contrast to the MHC genes found on Chr37, we found that the immunoglobulin new antigen receptor (IgNAR)^19^ family (four complete IgNAR structure, CH1-CH2-CH3-CH4-CH5-V, and two incomplete IgNAR) was located on Chr44 (**Fig. 1e**), again indicating the high impact of chromosome rearrangements on immune system evolution.

In addition to fast-evolving chromosomes and genes, chromosome rearrangements may also play an important role in gene loss events. Comparisons of chicken, zebrafish, medaka and bamboo shark showed at least four possible genome rearrangement events in the bamboo shark resulting in the loss of the gene, *p2rx5* (**Fig. 2a**), which was previously reported to be involved in bone development and homeostasis^20–26^. Furthermore, analysis of the whole gene family of purinergic receptor P2X in twelve species (including sea lamprey, the three sharks and eight bony fishes) showed that *p2rx3* and *p2rx5* were lost in the three cartilaginous fishes while at least five paralogs (*p2rx1, p2rx2, p2rx3, p2rx4, p2rx5*) with multiple copies were found in bony fishes (**Supplementary Figure 12** and **Supplementary Table 14-15**). P2X receptors consists of ligand-gated ion channels, and activation of the receptor triggers signaling pathways associated with Ca^2+^ influx^27–29^. Moreover, P2X receptors has been showed to participate in differentiation and proliferation of osteoblast^20,29,30^, bone formation and resorption^24,31,32^. Thus, it is reasonable to assume that loss of those genes during evolution may explain the establishment of chondrification of the endoskeleton in cartilaginous fish. To verify our hypothesis, we performed a knockout of *p2rx3a* and *p2rx5* using a CRISPR/Cas9 approach in zebrafish embryos **(Supplementary Figure 13)**, and the embryos of 9 days post-fertilization (dpf) were stained with alizarin red. Ventral view of 9 dpf mutant embryos showed significantly reduced mineralization in multiple skeletal landmarks(**Fig. 2b, c, d, f, g**), confirmed by semi-quantitative analysis of mineralization degree (mineralization area and mineralization integral optical density, IOD) in those embryos (**Fig. 2h, i, j, k**). Despite the extent of bone reduction was various due to somatic chimaerism with regard to gene disruption, the phenotype of inhibited mineralization took up a noticeable greater proportion in the knockout embryos, when compared to the wild type (**Fig. 2e**). These results in zebrafish suggest that the absence of these genes is one of the major causes of the chondrified endoskeleton in cartilaginous fish, underscoring the functional importance of genome rearrangements in evolution.

**Fig. 2.**
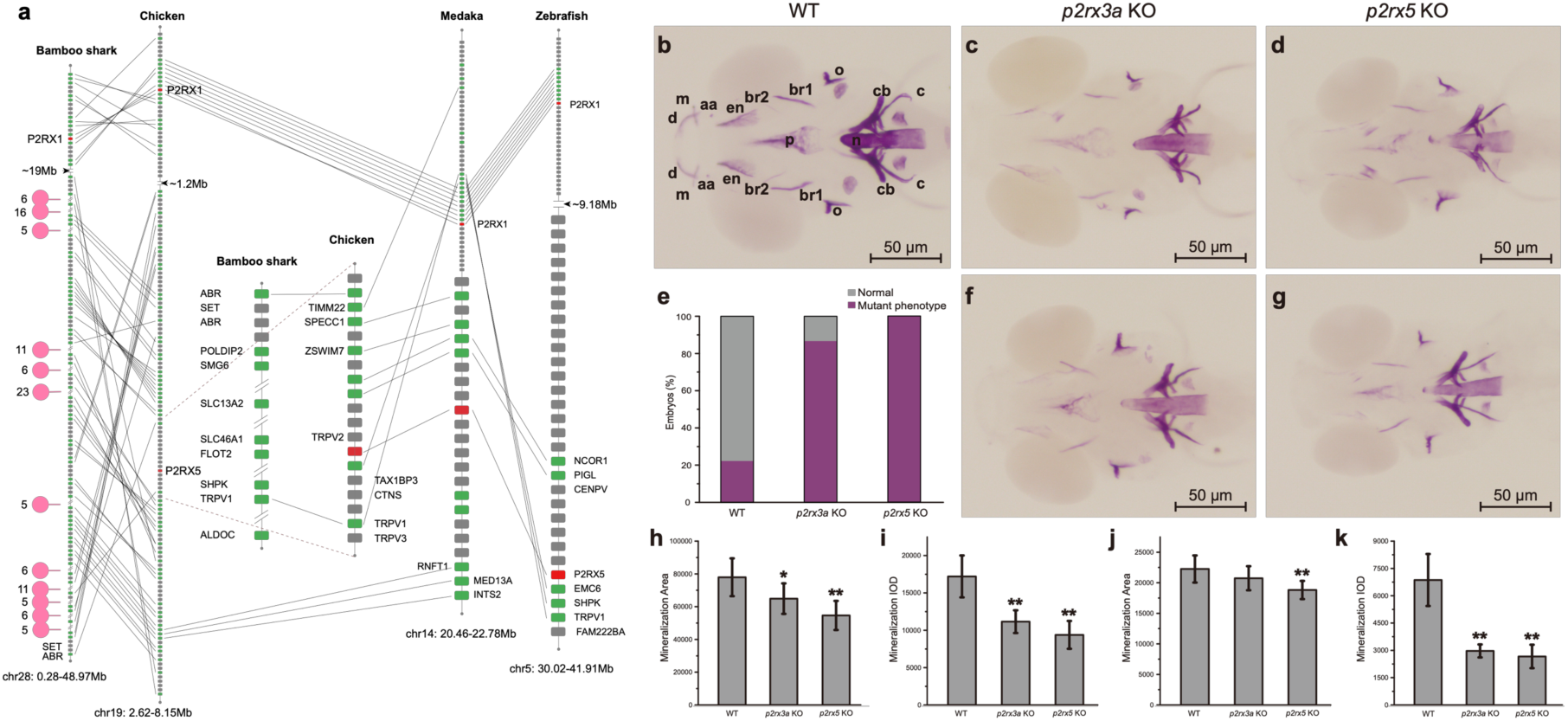
The loss of *p2rx5* gene in bamboo shark and targeted mutagenesis of zebrafish *p2rx3a* and *p2rx5* genes by CRISPR/Cas9 results in deformed or reduced bone development. **a)** The syntenic relationship of the *p2rx5* region among bamboo shark, chicken, medaka and zebrafish. The red rectangles represent P2X genes. The lines represent syntenic genes. The numbers besides the pink dots represent gene number in the gap regions. **b)** Alizarin red staining of 9-dpf wild type zebrafish embryo; the 11 landmarks are branchiostegal ray1 (br1), cleithrum (c), ceratobranchial 5 (cb), ceratohyal (ch), dentary (d), entopterygoid (en), hyomandibular (hm), maxilla (m), notochord (n), opercle (o), parasphenoid (p). **c, f)** Alizarin red staining of 9-dpf zebrafish embryo injected with Cas9 protein and single guide RNA (sgRNA) targeting exon 1 of the *p2rx3a* gene. Bone development was significantly reduced or deformed in multiple landmark positions. **d, g)** Alizarin red staining of 9-dpf zebrafish embryo injected with Cas9 protein and sgRNA targeting exon 1 of the *p2rx5* gene. Bone development was significantly deformed and strongly reduced in multiple landmark positions. **e)** Proportion of reduced or deformed bone phenotype in genotypic mutant embryos or wild type control. Phenotypic mutation rates were significantly higher in *p2rx3a* (15 of 15, 100%) and *p2rx5* (65 of 75, 86.67%) knocked out individuals than in wild type control (injected with phenol red, 12 of 54, 22.22%) (*P* < 0.01). **h, i)** The effect of gene knockout on head skeleton mineralization area and mineralization integral optical density (IOD) in 9-dpf zebrafish. A significant reduction of mineralization area and IOD were observed in both *p2rx3a* (*n* = 15) and *p2rx5* (*n* = 33) knocked out larvae, compared to wild type (*n* = 33)(\**P* < 0.05, \*\**P* < 0.01). **j, k)** The effect of gene knockout on notochord mineralization area and mineralization IOD in 9-dpf zebrafish. A significant reduction of mineralization IOD were observed in both *p2rx3a* (*n* = 15) and *p2rx5* (*n* = 33) knocked out larvae, compared to wild type (*n* = 33), and mineralization area was significantly shrunk in *p2rx5* knocked out larvae (\*\**P* < 0.01).

## Discussion

With whole genome and Hi-C sequencing, we successfully constructed a chromosome-level genome of the bamboo shark which is, to our knowledge, the first chromosome-level genome assembly for cartilaginous fish species. Guided by the previous observation of highly divergent chromosome numbers in cartilaginous fishes, we inferred the ancestral chromosomes of cartilaginous fishes to find dynamically rearranging chromosomes during their evolution. We then illustrated the evolutionary consequences of these rearrangements, including the formation of fast-evolving chromosomes with fewer conserved genes and more fast-evolving genes, as well as the loss of functionally important genes. We also inferred that these consequences ultimately resulted in phenotypic diversity, including the formation of immune systems specific to sharks and the emergence of cartilage. Our study highlights the importance of chromosome rearrangements in the diversification of cartilaginous fish, and also provides numerous insights into evolution of all species. With the highly effective methods described in this study, more chromosome-level genome assemblies can be constructed in the future to further illustrate the evolution of the oldest jawed vertebrate groups, as well as the paraphyletic group of all fishes. Our bamboo shark genome will also be significant for future immune studies and related pharmaceutical development.

## Materials and Methods

### DNA, RNA extraction

Blood samples were collected from a female bamboo shark for whole genome sequencing. 14 tissue types were collected for RNA sequencing. A DNA sample was extracted from whole blood using the phenol-chloroform method and its quality and quantity were assessed by pulsed field gel electrophoresis and Qubit Fluorometer. RNA samples were extracted by using TRIzol^®^ Reagent and was assessed by Agilent 2100.

### Library construction

Firstly, for WGS libraries with average insert sizes of 180 bp and 350 bp, a Covaris E220 ultrasonicator (Covaris, Brighton, UK) was used to shear the extracted high-quality DNA and AMPure XP beads (Agencourt, Beverly, the U.S.) were used to select target fragments. Then, fragment end-repairing and A-tailing were performed by T4 DNA polymerase, T4 polynucleotide kinase and rTaq DNA polymerase. Next, PCR amplification of eight cycles was carried out and the single-strand circularization process was performed using T4 DNA Ligase, generating a single-stranded circular DNA library for sequencing.

Secondly, for mate-pair libraries, a Covaris E220 was used to acquire ∼2 kb DNA fragments and Hydroshear (GeneMachines, CA, USA) was used to acquire ∼5 kb, ∼10 kb and ∼20 kb DNA fragments. After further selection and purification of DNA, fragments were end-repaired and biotin-labeled. The modified fragments were circularized and re-fragmented using a Covaris E220. Biotin-labeled DNA fragments were captured on M280 streptavidin beads (Invitrogen, CA, USA), end-repaired, A-tailed and adaptor-ligated. Biotin-labeled fragments were PCR-amplified, purified on streptavidin-coated magnetic beads, size-selected by agarose gel electrophoresis and column purification, single-stranded and re-circularized. The purified PCR products were heat-denatured together with an adapter that was a reverse-complement to a particular strand of the PCR product, and the single-stranded molecule was then ligated using DNA ligase to get a single-stranded circular DNA library.

Thirdly, a blood sample was used for constructing the Hi-C library. The fresh blood cells (5 × 106) were collected by centrifugation and re-suspended in 1 ml of 1x PBS by repetitive pipetting. The cells were cross-linked by adding 37% formaldehyde (SIGMA, America) to obtain a 1% final concentration, to which was added a 2.5M glycine solution (SIGMA, America) to a final concentration of 0.2M to quench the reaction. To prepare nuclei, the formaldehyde-fixed powder was resuspended in nuclei isolation buffer (10 mM Tris-HCl pH 8.0 (SIGMA, America), 10 mM NaCl (BEYOTIME, Shanghai, China), 1× PMSF (SIGMA, St. Louis, America)) and then incubated in 0.5% SDS for 10 min at 62 °C. SDS was immediately quenched with 10% Triton X-100 (SIGMA, St. Louis, America) and the nuclei were collected by brief centrifugation. DNA was digested by restriction enzyme (Mbo I) (NEB, Ipswich, America) and the 5’ overhang was repaired using a biotinylated residue (0.4 mM biotin-14-Datp (INVITROGEN, America). The resulting blunt-end fragments were ligated in situ (10X NEB T4 DNA ligase buffer (NEB, Ipswich, America), 10% Triton X-100 (SIGMA, St. Louis, America), 10 mg/ml BSA (NEB, Ipswich, America), T4 DNA ligase (NEB, Ipswich, America)). Finally, the isolated DNA was reverse-crosslinked (adding 10 mg/ml proteinase K (NEB, Ipswich, America) and 1% SDS (AMBION, Waltham, America) to the tube followed by incubation at 56°C overnight) and purified (by putting the reverse-crosslinked DNA liquid into three tubes equally, adding 1.5× volumes of AMpure XP (AGENCOURT, Brea, America) mixture to each tube, vortexing and spinning down briefly, incubating for 10 min. at room temperature, placing on the MPS (INVITROGEN, Waltham, America) for 5 min. at room temperature, discarding supernatant, washing the beads twice with 1 ml of freshly made 75% ethanol (SINOPHARM, Shanghai, China), air-drying the beads completely and resuspending the beads in 30 µl of ddH2O). The Hi-C library was created by shearing 20 ug of DNA and capturing the biotin-containing fragments on streptavidin-coated beads using Dynabeads MyOne Streptavidin T1 (INVITROGEN, Waltham, America). Then DNA fragments were end-repaired and adaptor ligation was performed. After PCR (95°C 3 min.; [98°C 20 sec., 60°C 15 sec., 72°C 15 sec.] (8 cycles); 72°C 10 min.), the standard circularization step required for the BGISEQ-500 platform was carried out and DNBs were prepared as previously described.

Fourthly, for RNA library construction, mRNA was extracted from different tissues using TRIzol^®^ Reagent, fragmented, and then reverse-transcribed into cDNA by using Hiscript II Reverse Transcriptase (Vazyme Biotech, Nanjing City, P.R. China). Then, all single-stranded circular DNA libraries were constructed by using the same strategy described as above.

### Sequencing for all libraries

All sequencing data were generated using the BGISEQ-500 platform. Libraries with an average insert size of 180 bp and 350 bp were sequenced yielding paired-end reads with 100 bp in length. Mate-pair libraries with average insert sizes of 2k, 5k, 10k and 20k and Hi-C library were sequenced yielding reads with 50 bp in length. RNA libraries were also sequenced yielding paired-end reads with 100 bp in length.

### Genome assembly and annotation

Firstly, we filtered raw sequencing data by discarding low-quality reads (defined as >10% bases with quality values less than 10 and >5% unidentified (N) bases), adaptor-contaminated reads and PCR duplicate reads. We trimmed a few bases at the start and end of reads according to the FastQC (v0.11.2)^33^ results. Secondly, we randomly selected about 40X clean reads to carry out k-mer analysis to estimate the genome size (**Supplementary Figure 1**). Thirdly, we used Platanus (v1.2.4)^34^ to perform the initial assembly with WGS clean data with parameters “assemble –k 29 –u 0.2, scaffold -l 3 -u 0.2 -v 32 -s 32 and gap_close –s 34 – k 32 –d 5000”. We filled gaps using KGF (v1.19) and GapCloser with default parameters. Fourthly, to obtain a chromosome-level genome, HIC-Pro^35^ was used for quality control of Hi-C sequencing data with parameters [BOWTIE2_GLOBAL_OPTIONS = --very-sensitive - L 30 --score-min L,-0.6,-0.2 --end-to-end –reorder;BOWTIE2_LOCAL_OPTIONS = --very-sensitive -L 20 --score-min L,-0.6,-0.2 --end-to-end –reorder; IGATION_SITE = GATC; MIN_FRAG_SIZE = 100; MAX_FRAG_SIZE = 100000; MIN_INSERT_SIZE = 50; MAX_INSERT_SIZE = 1500]. Finally, the software packages Juicer^36^ and 3d-dna^37^ were employed to generate contact matrices of chromatin and constructed chromosomes with parameter [-m haploid -s 4 -c 5].

After obtaining the final chromosome-level genome, we proceeded with genome annotation including repeat contents, gene models and gene function. For repeat section, both homolog-based and *de novo* prediction strategies were carried out. In detail, RepeatMasker (v 4.0.5)^38^ and RepeatProteinMasker (v 4.0.5) were used to detect interspersed repeats and low complexity sequences against the Repbase database^39^ at the nuclear and protein levels, respectively. Then RepeatMasker was further used to detect species-specific repeat elements using a custom database generated by RepeatModeler (v1.0.8) and LTR-FINDER (v1.0.6)^40^. In addition, Tandem Repeat Finder (v4.0.7) ^41^ was dispatched to predict tandem repeats. The final repeat content result was integrated using in-house scripts. Before gene model construction, we masked the repeat sequences because of their negative effect on gene model prediction. We downloaded protein sets of 13 species including *Homo sapiens, Mus musculus, Gallus gallus, Xenopus tropicalis, Ornithorhynchus anatinus, Danio rerio, Oryzias latipes, Strongylocentrotus purpuratus, Ciona intestinalis, Rhincodon typus* and *Callorhinchus milii* from RefSeq (release 82), *Petromyzon marinus* from Ensembl (release 84) and *Branchiostoma floridae* from JGI Genome Portal (http://genome.jgi-psf.org/Brafl1/Brafl1.home.html) and aligned them to the masked genome with BLAT^42^ to identify positive match regions. GeneWise (v2.2.0)^43^ was then used to do accurate alignments for target regions and to predict homolog-based gene models. Transcriptome reads from 14 tissues including blood, eye, gill, heart, liver, muscle, spleen, stomach, dorsal fin, tail fin, pancreas, leptospira, 2 capsulogenous gland and 2 kidney samples were mapped to the genome with HISAT2^44^ and StringTie^44^ was used to assemble gene transcripts. TransDecoder^45^ was then used to predict the candidate complete ORFs. Further, for *de novo* gene prediction, we employed AUGUSTUS (v3.1)^46^ to scan the whole genome with a custom training set generated by using 2,000 high quality genes. Subsequently, we combined the homology-based and *de novo*-predicted gene sets using GLEAN ^47^ and integrated the GLEAN and transcriptome results with in-house scripts to generate a representative and non-redundant gene set. The final gene set was assigned with a potential function by aligning proteins to databases including KEGG, Swissprot, TrEMBL and InterPro.

### Evolution of chromosomes

For ancestral chromosomes construction, we identified paralogous genes and orthologous genes by using criteria defined by *Salse et al*^9^ with both Cumulative Identity Percentage (CIP) and Cumulative Alignment Length Percentage (CALP) value of 0.5 and selected genes pairs defined as A match B best and B match A best. Then MCSCAN^48^ was used to generate synteny blocks with default parameters. We first noted 54 shared duplications (with 414 paralogous gene pairs) on all the chromosomes. After further integration of these duplications and gene pairs, we found 16 duplicated chromosome pairs and 5 single chromosomes. For identification of conserved genes among elephant shark, whale shark, spot gar, and medaka, the same criteria with both CIP and CALP value of 0.3 were used. The best hit of multiple matches was selected. Ks values of single copy genes were calculated by KaKs Calculator with default parameter. Heterozygosity of each chromosome was calculated by calling heterozygous SNPs generated by BWA^49^ and SAMtools package^50^.

### MHC genes and P2X gene family analysis

We downloaded coding sequences and proteins of *Callorhinchus milii* (GCF_000165045.1), *Rhincodon typus* (GCF_001642345.1), *Fugu rubripes* (GCF_000180615.1) and *Larimichthys crocea* (GCF_000972845.1) from NCBI database, *Danio rerio, Latimeria chalumnae, Oryzias latipes* and *Gasterosteus aculeatus* from Ensembl (release 84), sea lamprey from (https://genomes.stowers.org/organism/Petromyzon/marinus) and *Branchiostoma floridae* from JGI Genome Portal (http://genome.jgi-psf.org/Brafl1/Brafl1.home.html) and then performed initial filtering by discarding sequences less than 30 amino acids keeping the longest transcript if one gene was present with multiple transcripts. Next, the gene function annotation of these gene sets was performed by using the same criteria as bamboo shark. Then we summarized the MHC genes and P2X gene families by using the above information. The P2X-like genes (**Supplementary Table 14**) were searched in NCBI database.

### Identification of IgNAR

We downloaded IgNAR (immunoglobin new antigen receptor from cartilaginous) nucleotide sequences from NCBI database and align these sequences to bamboo shark genome using BLAST to verify the IgNAR loci.

### Knockout of P2X genes

CRISPR/Cas9 were used to cleave targeted sites of genes of interest in order to induce mutations during non-homologous end joining (NHEJ) repair in zebrafish. For *p2rx3a* and *p2rx5* genes, 3 different sgRNAs were designed to target exon 1 of *p2rx3a* and of *p2rx5* (**Supplementary Table 16**). sgRNAs were designed using software developed by an online tool of CRISPOR (http://crispor.tefor.net), and then synthesized and purified in our labs. Cas9 protein was bought from System Biosciences (2438 Embarcadero Way, Palo Alto, CA 94303). CRISPR/Cas9 molecules were transferred into one-cell staged zebrafish embryos by microinjection, 1 nL per injection with 200 pg nL^-1^ sgRNA and 375 pg nL^-1^ Cas9 protein. Tail clips of 9 dpf embryos were used for preparing genomic DNA, KOD-FX (TOYOBO, Japan) were employed for genotyping PCRs with primer pairs P2RX3F/R and P2RX5F/R (**Supplementary Table 17**) for *p2rx3a* and *p2rx5* assays, respectively. Purified PCR products were cloned into pMD18-T vector (TaKaRa, Japan) for transformation to DH5-α competence cells of *Escherichia coli* and sequencing.

Embryos were then stained with alizarin red that binds to calcium salt in mineralization matrix for phenotyping. Alizarin red staining followed a modified protocol^51^. Phenotype data were collected by microfilm photos, and all images for each test were taken under identical conditions. The mineralization area and integral optical density (IOD) of alizarin red staining were quantified with the software of Image-Pro Plus V6.0 (Media Cybernetics, USA).

## Acknowledgments

We would like thank Dr. Lynn Fink for revising these manuscripts here. This work was supported by Shenzhen-Hongkong Collaboration Fund JCYJ20170412152916724 (20170331), State Key Laboratory of Agricultural Genomics (No. 2011DQ782025) and National Key R&D Program of China (2018YFD0900301).

## Author contributions

X.L., N.Y., G.F. and X.X. designed and managed this project. M.W., C.L., H.X., L.W., H.R., Y.X., Q.X., and S.P., were responsible for collecting samples, library construction, sequencing and co-drafting the manuscript. Y.Z., H.G., and J.G., worked on genome assembly, annotation, chromosome evolution, transcriptome and co-drafting the manuscript. J.W., M.L., X.G., Q.L. Y.S., and Y.L., performed data processing WGD, gene family analysis and repeat analysis. H.L., Y.G., Q.Z., C.W., L.S., X.D. carried out CRISPR/Cas9 experiments. J.S., S.L., and S.L helped to revise the manuscripts. All authors took part in the interpretation of data.

## Competing interests

The authors declare that they have no competing interests.

## Data and materials availability

This Whole Genome Shotgun project has been deposited at DDBJ/ENA/GenBank under the accession QPFF00000000 referring project PRJNA478295. Raw RNA sequencing reads have been also uploaded to the SRA database under accession SRP154403. The assembled genome can also be obtained from CNSA (CNGB Nucleotide Sequence Archive) by assembly ID: CNA0000025.

