## Supplementary Figures and Tables for "Dynamic chromosome rearrangements of the white-spotted bamboo shark shed light on cartilaginous fish diversification mechanisms"

[1]BGI-Qingdao, BGI-Shenzhen, Qingdao 266555, China
[2]BGI-Shenzhen, Shenzhen 518083, China
[3]State Key Laboratory of Quality Research in Chinese Medicine and Institute of Chinese Medical Sciences, Macao, China
[4]China National GeneBank, BGI-Shenzhen, Shenzhen 518120, China
[5]City University of Hongkong, Kowloon, Hongkong SAR
[6]Complete Genomics, Inc., San Jose, CA 95134, USA
[7]Yellow Sea Fisheries Research Institute, CAFS, Key Laboratory for Sustainable Development of Marine Fisheries, Ministry of Agriculture, Qingdao 266071, China
[8]James D. Watson Institute of Genome Sciences, Hangzhou 310058, China
[9]Department of Biological Sciences, King Abdulaziz University (KAU), Jeddah 21589, Saudi Arabia
[†]These authors contributed equally to this work.

### Supplementary Figures

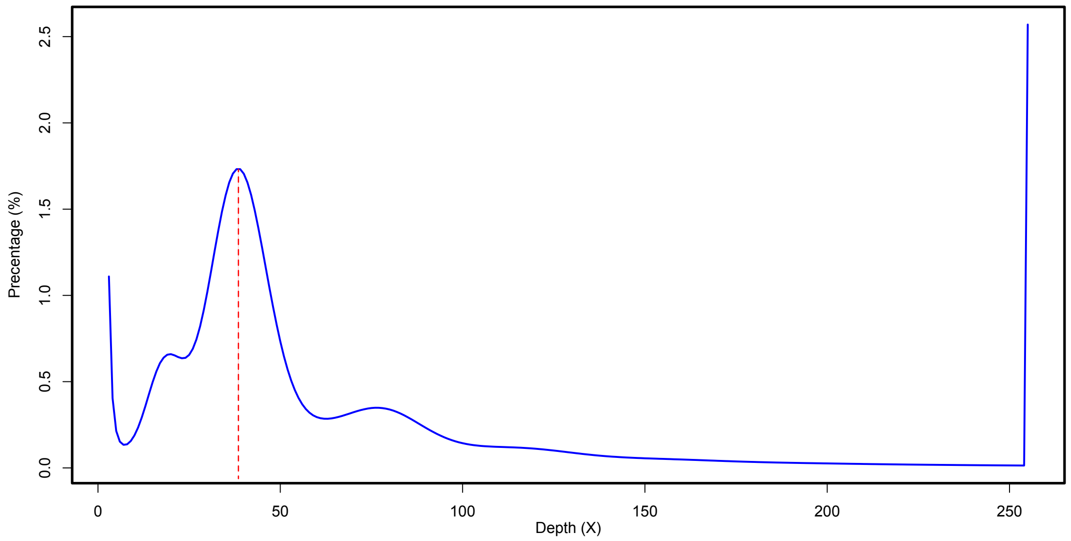

**Supplementary Figure 1.** 17-mer frequency distribution of the bamboo shark genome. A total of 209 Gb sequencing data was used for this k-mer analysis, generating 169,452,206,642 k-mers. The red dotted line represents the main peak-depth value, around 39. The estimated genome size (total num of kmer/peak-depth) is about 4.30Gb. From this plot, we can easily identify heterozygous and duplicated peaks which indicate this genome is complex.

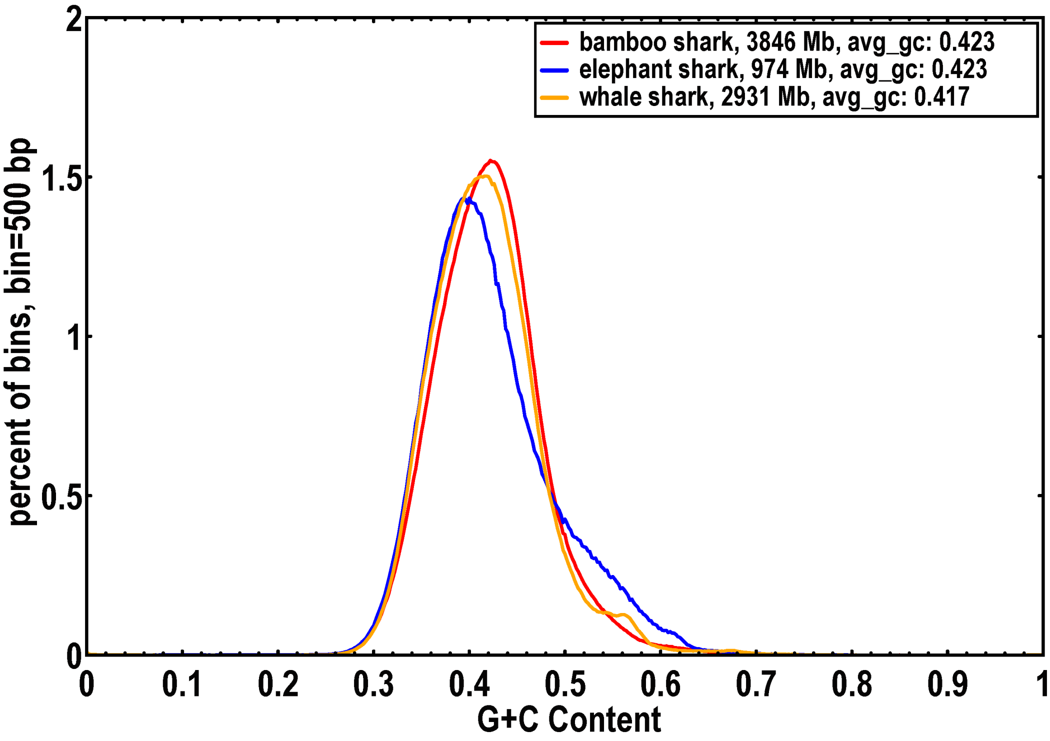

**Supplementary Figure 2.** GC content of the assembled bamboo shark, elephant shark, and whale shark genomes.

GC-content (*x*-axis) is quantified as the final proportion of 500 bp windows or bins (*y*-axis).

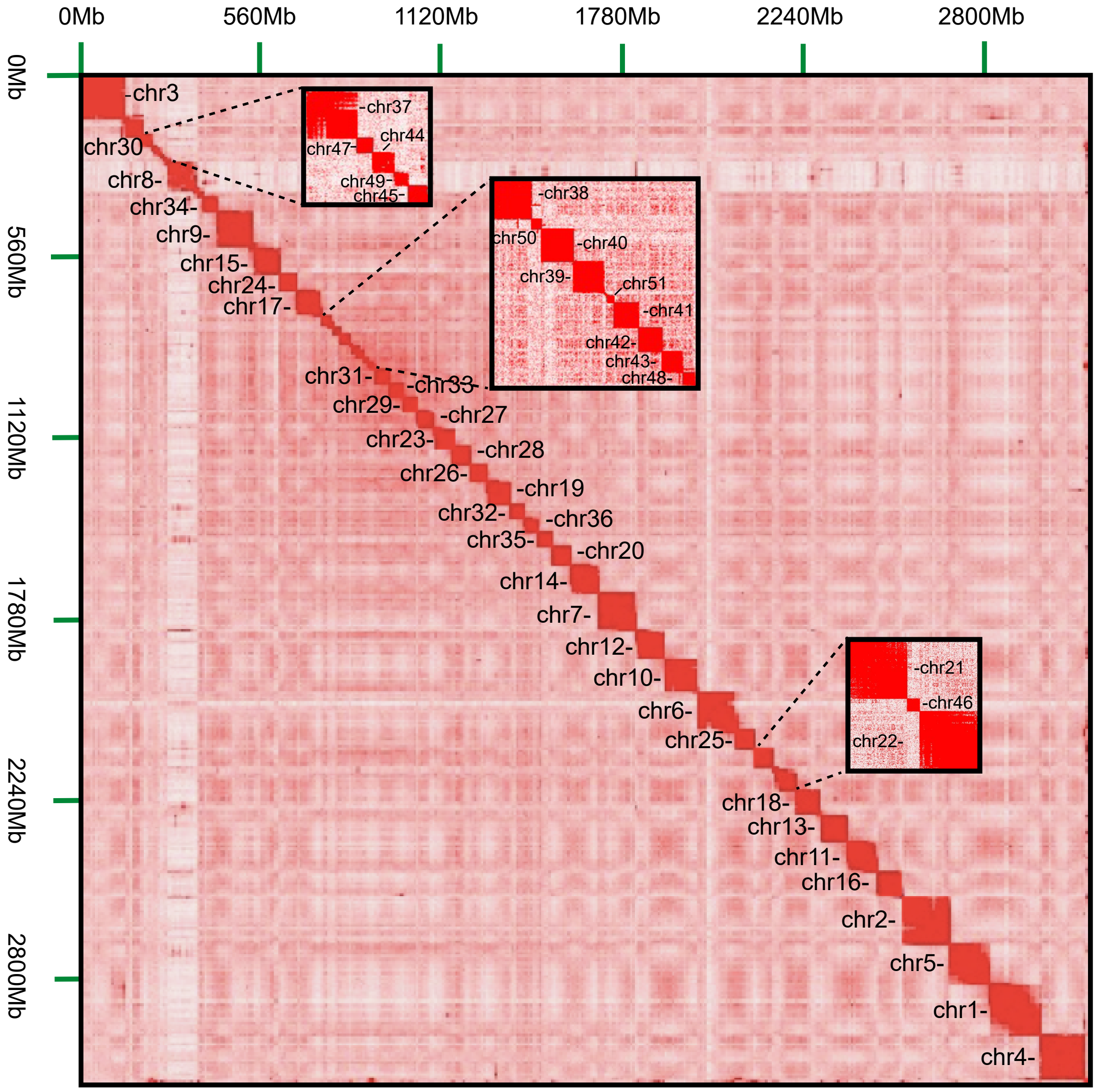

**Supplementary Figure 3.** Heat map of chromatin interaction relationships at 125 kb resolution of 51 chromosomes.

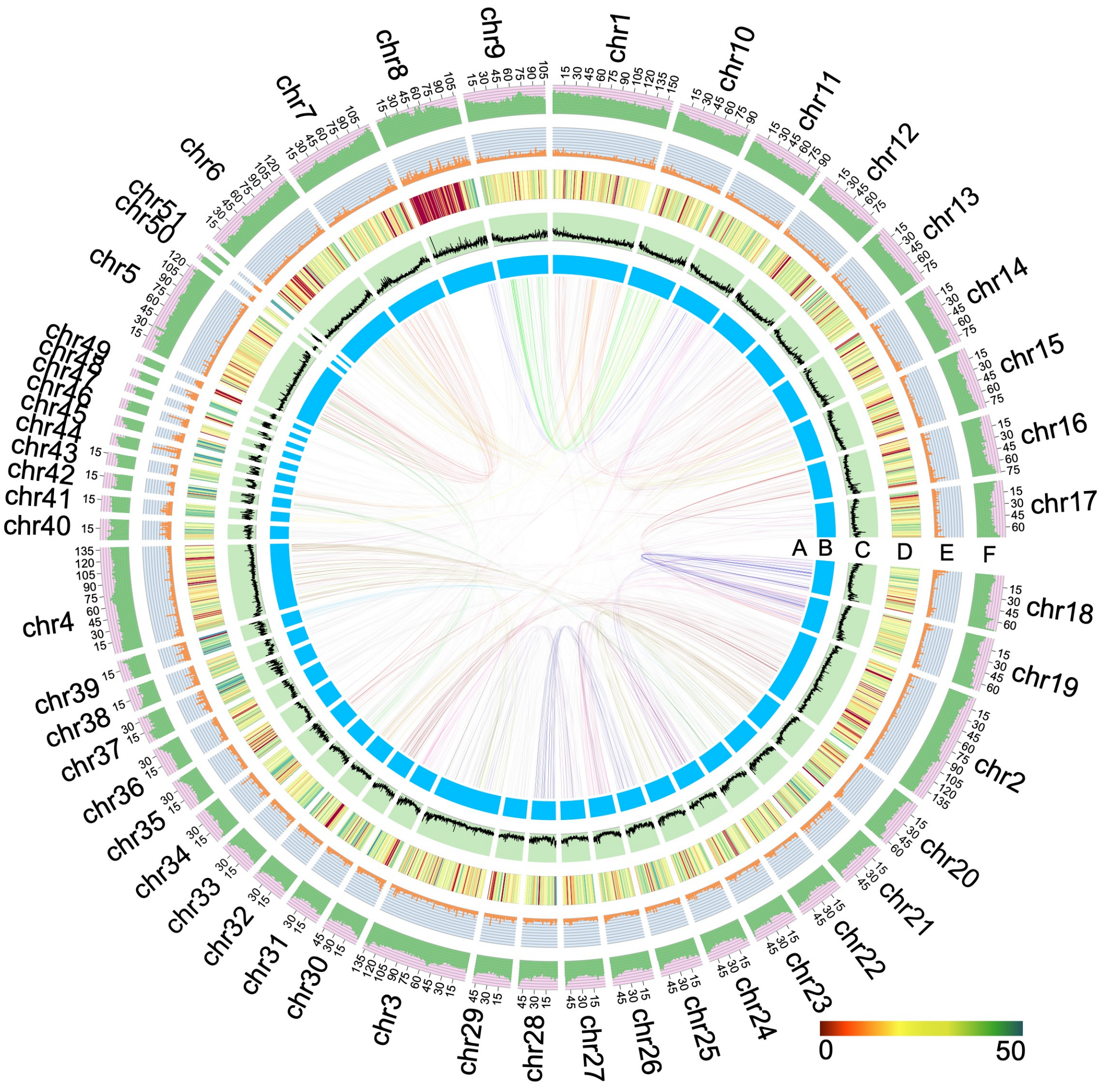

**Supplementary Figure 4.** Circos plot of the genomic landscape.

A, Paralogous gene pairs of 51 chromosomes. B, 51 chromosomes of the assembled genome. C, GC content of the assembled genome at 100kb windows. D, gene density per Mb on each chromosome. E, histogram of DNA transposon ratios. F, histogram of retrotransposon ratios.

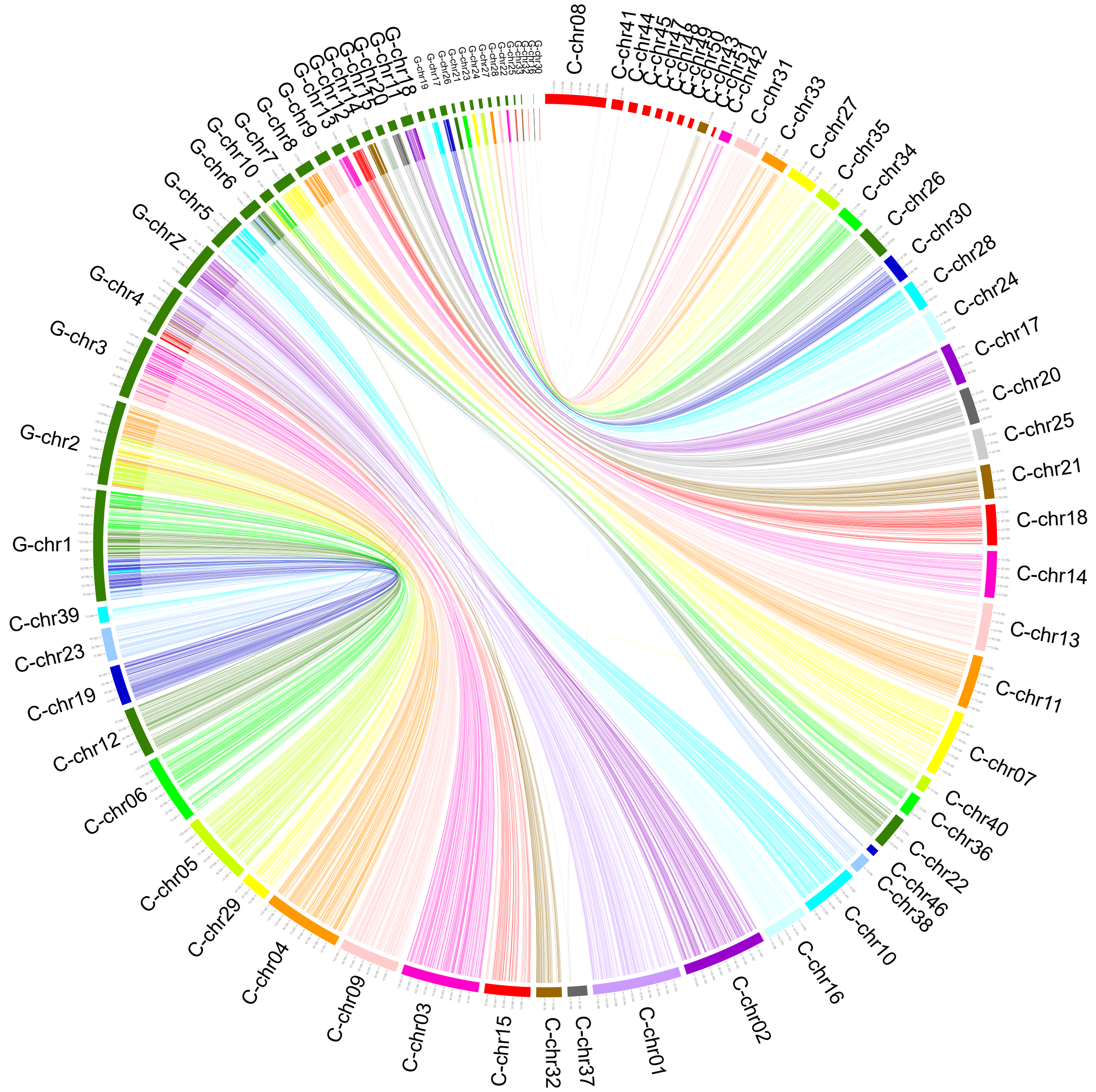

**Supplementary Figure 5.** Circos plot of comparisons between bamboo shark and chicken.

G-chr and C-chr represent chicken and bamboo shark chromosomes, respectively. The lines represent gene pairs identified with both CIP and CALP (defined by *Salse et al (1)*) value of 0.3.

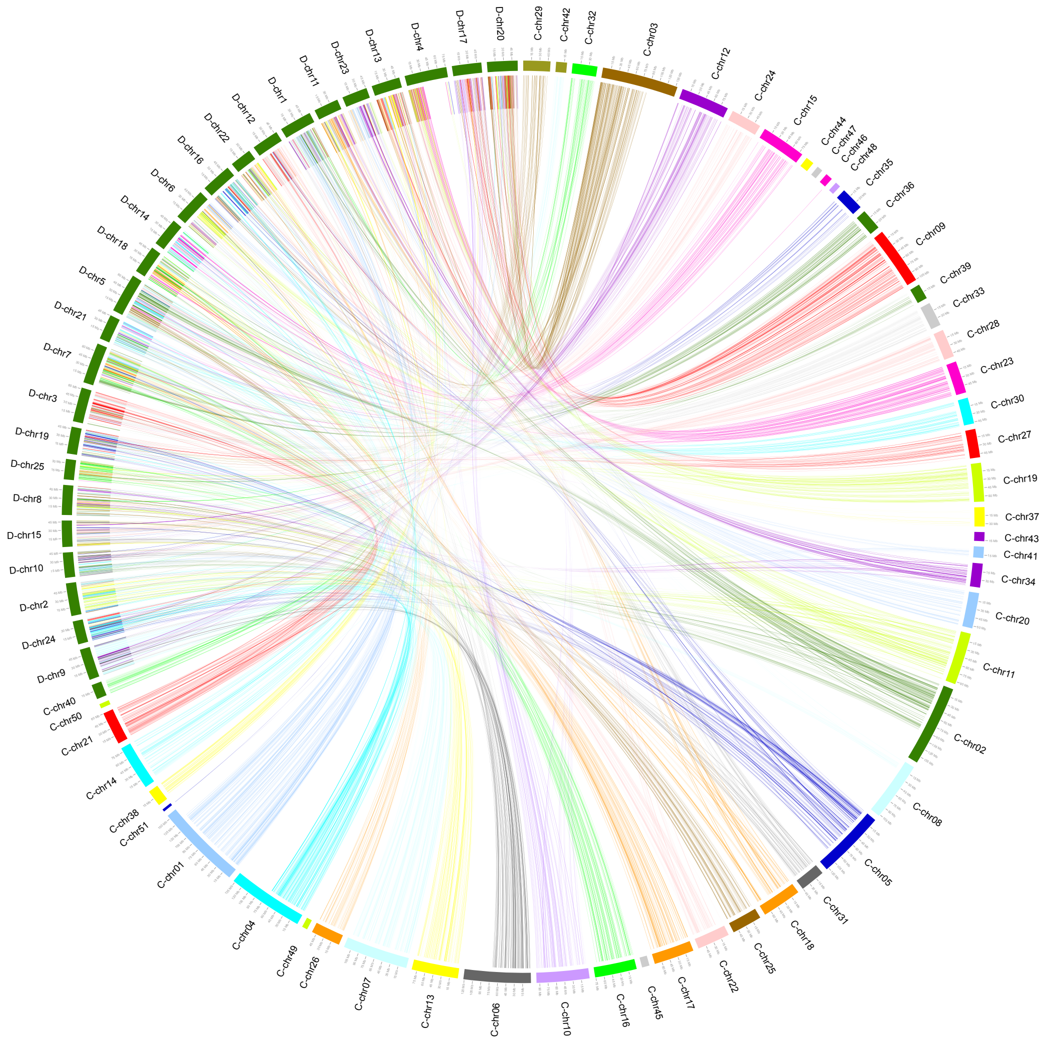

**Supplementary Figure 6.** Synteny relationship of bamboo shark and zebrafish.

D-chr and C-chr represent zebrafish and bamboo shark chromosomes, respectively. The lines represent gene pairs identified with both CIP and CALP (defined by *Salse et al (1)*) value of 0.3.

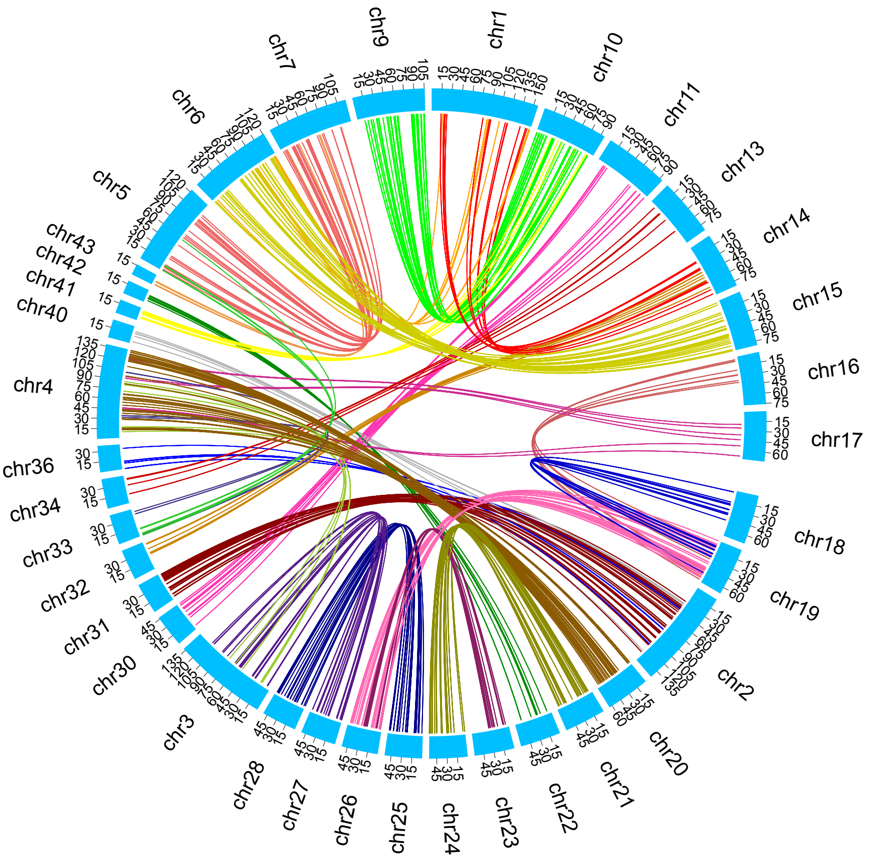

**Supplementary Figure 7.** Distribution of shared paralogous genes of bamboo shark and elephant shark. The lines represent gene pairs obtained by using blastp with both CIP and CALP (defined by *Salse et al(1)*) value of 0.5.

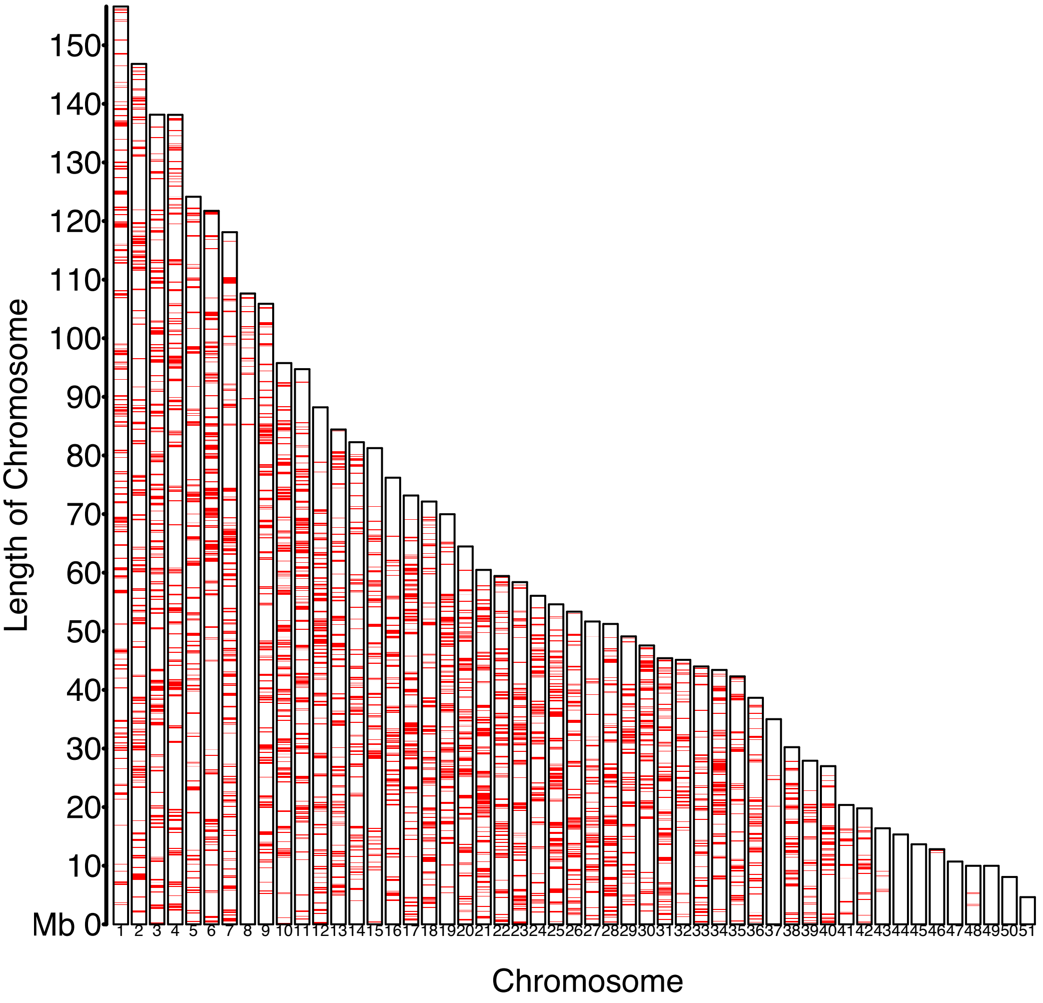

**Supplementary Figure 8.** Distributions of conserved regions of three cartilaginous fishes on 51 chromosomes of the bamboo shark.

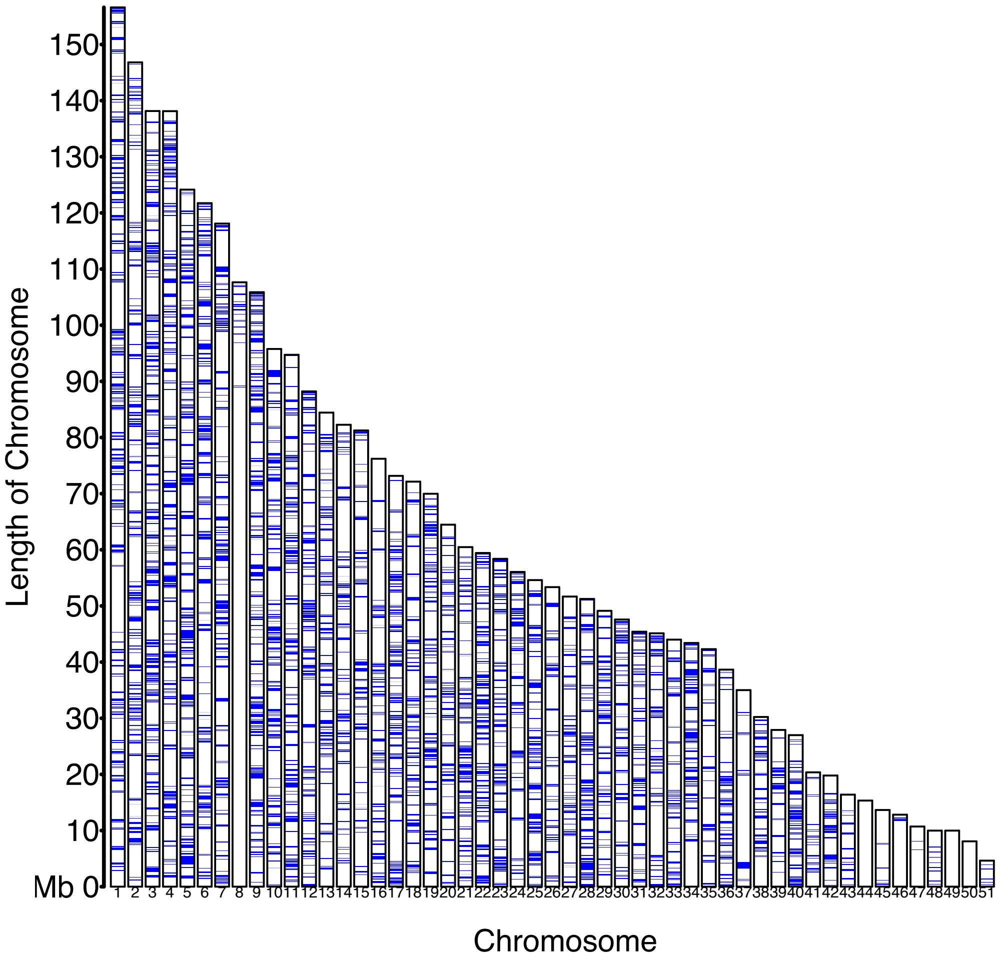

**Supplementary Figure 9.** Distributions of conserved regions between bamboo shark and medaka on 51 chromosomes of the bamboo shark.

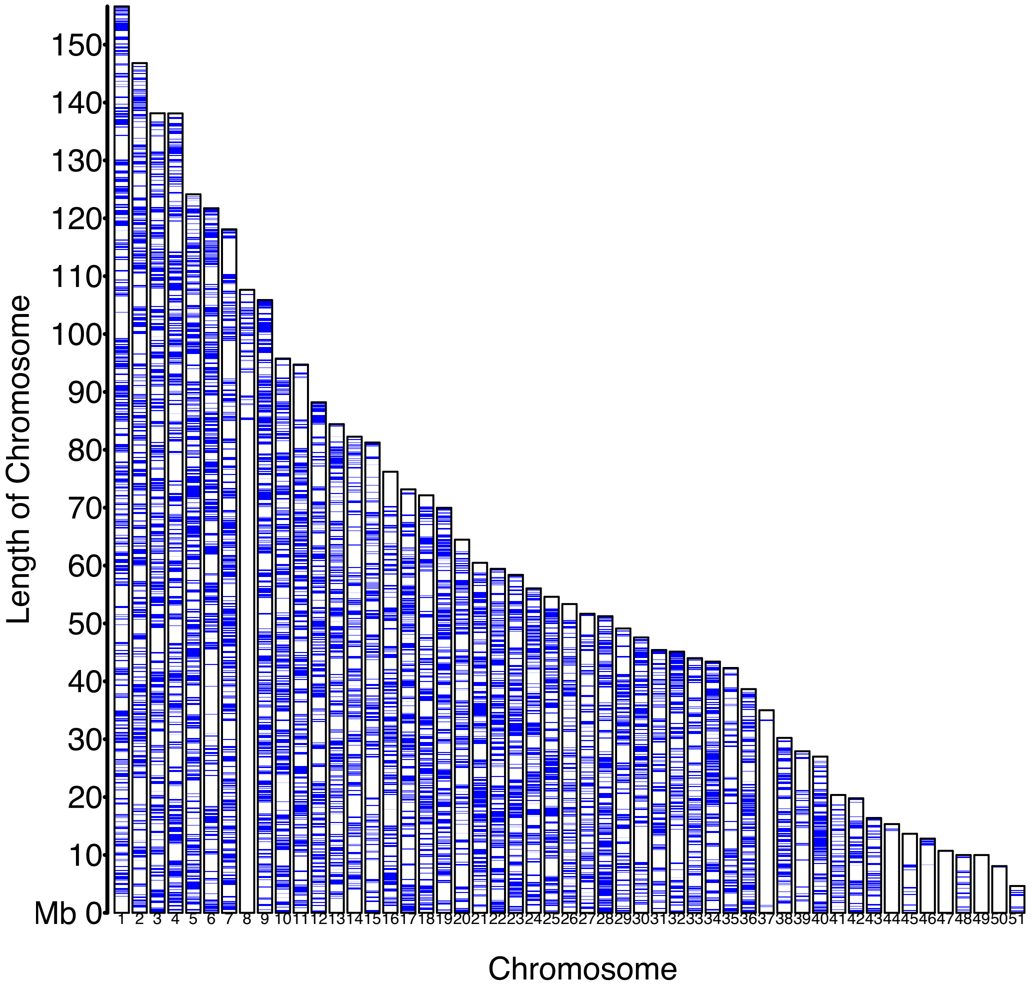

**Supplementary Figure 10.** Distributions of conserved regions between bamboo shark and spotted gar on 51 chromosomes of the bamboo shark.

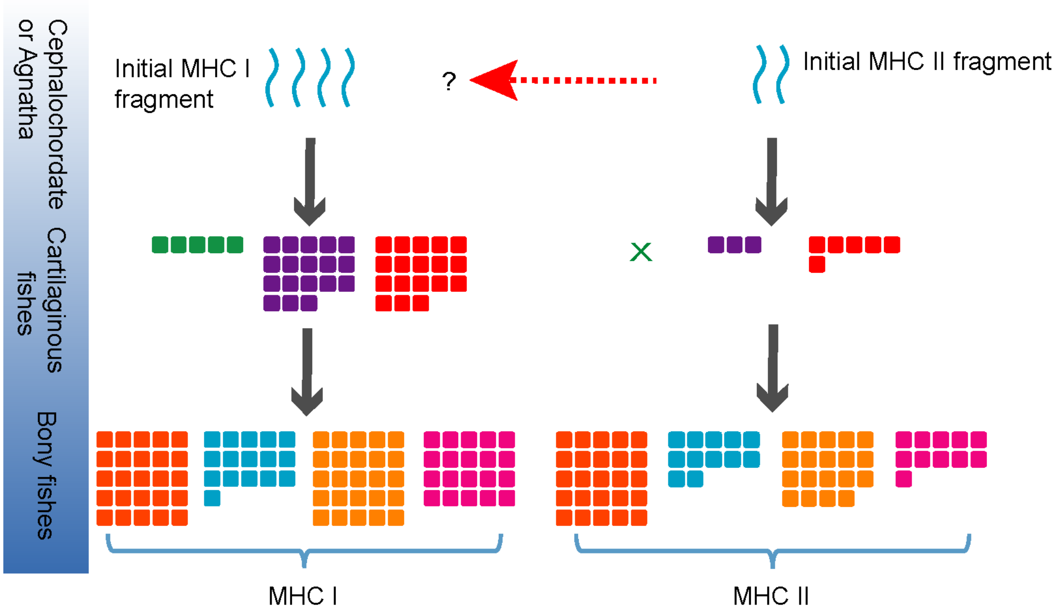

**Supplementary Figure 11.** Hypothetical model of MHC genes evolution.

The square numbers represent gene numbers. Green, elephant shark. Purple, whale shark. Red, bamboo shark. Orange, zebrafish. Blue, medaka. Yellow, coelacanth, magenta, three-spined sticklebacks. The green X mark represents there are no MHC class II genes found in the elephant shark genome. The red arrow and question mark represent the hypothesis that MHC class I molecules may be derived from MHC class II while more researches should be done in the future.

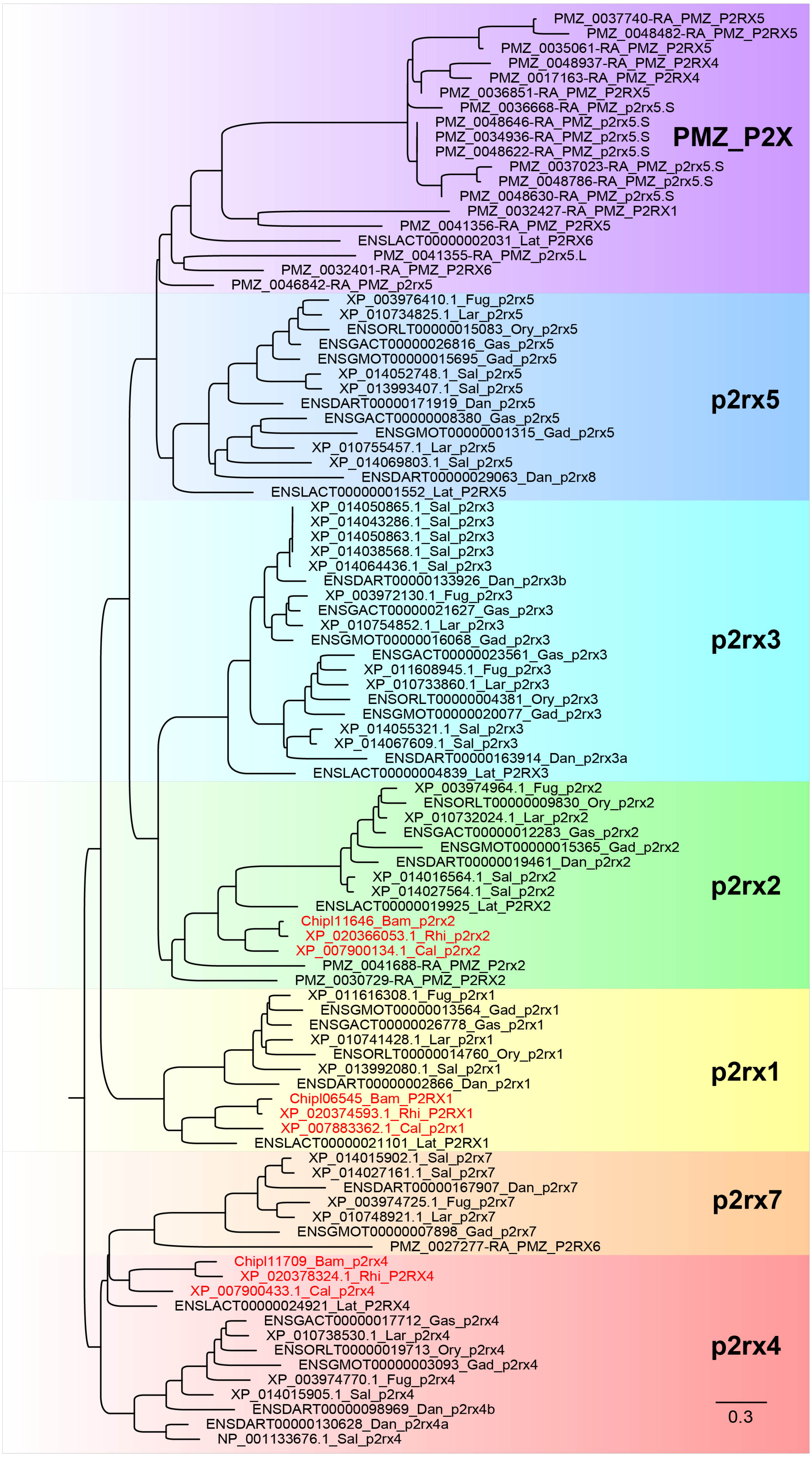
PMZ_0037740-RA_PMZ_P2RX5
PMZ_0048482-RA_PMZ_P2RX5
PMZ_0035061-RA_PMZ_P2RX5
PMZ_0048937-RA_PMZ_P2RX4
PMZ_0017163-RA_PMZ_P2RX4
PMZ_0036851-RA_PMZ_P2RX5
PMZ_0036668-RA_PMZ_p2rx5.S
PMZ_0048646-RA_PMZ_p2rx5.S
PMZ_0034936-RA_PMZ_p2rx5.S
PMZ_0048622-RA_PMZ_p2rx5.S
PMZ_0037023-RA_PMZ_p2rx5.S
PMZ_0048786-RA_PMZ_p2rx5.S
PMZ_0048630-RA_PMZ_p2rx5.S
PMZ_0032427-RA_PMZ_P2RX1
PMZ_0041356-RA_PMZ_P2RX5
ENSLACT00000002031_Lat_P2RX6
PMZ_0041355-RA_PMZ_p2rx5.L
PMZ_0032401-RA_PMZ_P2RX6
PMZ_0046842-RA_PMZ_p2rx5
PMZ_P2X
XP_003976410.1_Fug_p2rx5
XP_010734825.1_Lar_p2rx5
ENSORLT00000015083_Ory_p2rx5
ENSGACT00000026816_Gas_p2rx5
ENSGMOT00000015695_Gad_p2rx5
XP_014052748.1_Sal_p2rx5
XP_013993407.1_Sal_p2rx5
ENSDART00000171919_Dan_p2rx5
ENSGACT00000008380_Gas_p2rx5
ENSGMOT00000001315_Gad_p2rx5
XP_010755457.1_Lar_p2rx5
XP_014069803.1_Sal_p2rx5
ENSDART00000029063_Dan_p2rx8
ENSLACT00000001552_Lat_P2RX5
p2rx5
XP_014050865.1_Sal_p2rx3
XP_014043286.1_Sal_p2rx3
XP_014050863.1_Sal_p2rx3
XP_014038568.1_Sal_p2rx3
XP_014064436.1_Sal_p2rx3
ENSDART00000133926_Dan_p2rx3b
XP_003972130.1_Fug_p2rx3
ENSGACT00000021627_Gas_p2rx3
XP_010754852.1_Lar_p2rx3
ENSGMOT00000016068_Gad_p2rx3
ENSGACT00000023561_Gas_p2rx3
XP_011608945.1_Fug_p2rx3
XP_010733860.1_Lar_p2rx3
ENSORLT00000004381_Ory_p2rx3
ENSGMOT00000020077_Gad_p2rx3
XP_014055321.1_Sal_p2rx3
XP_014067609.1_Sal_p2rx3
ENSDART00000163914_Dan_p2rx3a
ENSLACT00000004839_Lat_P2RX3
p2rx3
XP_003974964.1_Fug_p2rx2
ENSORLT00000009830_Ory_p2rx2
XP_010732024.1_Lar_p2rx2
ENSGACT00000012283_Gas_p2rx2
ENSGMOT00000015365_Gad_p2rx2
ENSDART00000019461_Dan_p2rx2
XP_014016564.1_Sal_p2rx2
XP_014027564.1_Sal_p2rx2
ENSLACT00000019925_Lat_P2RX2
Chipl11646_Bam_p2rx2
XP_020366053.1_Rhi_p2rx2
XP_007900134.1_Cal_p2rx2
PMZ_0041688-RA_PMZ_P2rx2
PMZ_0030729-RA_PMZ_P2RX2
p2rx2
XP_011616308.1_Fug_p2rx1
ENSGMOT00000013564_Gad_p2rx1
ENSGACT00000026778_Gas_p2rx1
XP_010741428.1_Lar_p2rx1
ENSORLT00000014760_Ory_p2rx1
XP_013992080.1_Sal_p2rx1
ENSDART00000002866_Dan_p2rx1
Chipl06545_Bam_P2RX1
XP_020374593.1_Rhi_P2RX1
XP_007883362.1_Cal_p2rx1
ENSLACT00000021101_Lat_P2RX1
p2rx1
XP_014015902.1_Sal_p2rx7
XP_014027161.1_Sal_p2rx7
ENSDART00000167907_Dan_p2rx7
XP_003974725.1_Fug_p2rx7
XP_010748921.1_Lar_p2rx7
ENSGMOT00000007898_Gad_p2rx7
PMZ_0027277-RA_PMZ_P2RX6
p2rx7
Chipl11709_Bam_p2rx4
XP_020378324.1_Rhi_P2RX4
XP_007900433.1_Cal_p2rx4
ENSLACT00000024921_Lat_P2RX4
ENSGACT00000017712_Gas_p2rx4
XP_010738530.1_Lar_p2rx4
ENSORLT00000019713_Ory_p2rx4
ENSGMOT00000003093_Gad_p2rx4
XP_003974770.1_Fug_p2rx4
XP_014015905.1_Sal_p2rx4
ENSDART00000098969_Dan_p2rx4b
ENSDART00000130628_Dan_p2rx4a
NP_001133676.1_Sal_p2rx4
p2rx4
0.3

**Supplementary Figure 12.** Phylogenetic tree of the P2X gene family in 12 species.

Dan: zebrafish, Gad: atlantic cod, Gas: three-spined sticklebacks, Ory: medada, Lat: coelacanth, Sal: atlantic salmon, Fug: torafugu, Lar: large yellow croaker, Rhi: whale shark, Cal: elephant shark, Bam: bamboo shark, PMZ: sea lamprey. The shark's P2X genes are marked red. Although many *p2rx5* genes exist in PMZ, all of these genes were less than 246 aa in length (10 out of 14 less than 200aa). This tree was constructed with protein sequences of these genes by using softwares of MUSCLE (v3.8.31) and FastTree (version 2.1.10)

```
p2rx3a_Target_site_1
WT     TGAAGAGCTGGTCTGTTGGAATAATCAACCGGGTCGTTCAGCTGTTAATTATTTT←307 bp→GTTAACCTAATTAATCTAGCAAA
M1.1   TGAAGAGCTGGTCTGTTCAGCTGC....................TAATTATTTT  (-28,+7)
M1.2   TGAAGAGCTGCTACCTAAACGAAGAGCT←334 bp→GTTAACCTAATTAATCTAGCAAA  (-352,+18)
M1.3   TGAAGAGCTGGTCTGTTGGAATAATC...CGGGTCGTTCAGCT  (-3,+0)
M1.4   TGAAGAGCTGGTCTGTTGGAATAATC....GGGTCGTTCAGCT  (-4,+0)
M1.5   TGAAGAGCTGGTCTGTTGGAATAATCATCCGGGTCGTTCAGCT  (-1,+1)
M1.6   TGAAGAGCTGGTCTGTTGGAATAATCAAACCGGGTCGTTCAGCT  (-0,+1)
M1.7   TGAAGAGCTGGTCTGTTGGAATAATAATAACCGGGTCGTTCAGCT  (-1,+3)
M1.8   TGAAGAGCTGGTCTGTTGGAATAATCACCGGGTCATAATACCGGGTCGTTCAGCT  (-0,+12)
M2.1   TGAAGAG.....................CGGGTCGTTCAGCT  (-22,+0)
M2.2   TGAAGAGCTGGTCTGTTGGAATAA....CCGGGTCGTTCAGCT  (-4,+0)
M2.3   TGAAGAGCTGGTCTGTTGGAAT....CGTCGGGTCGTTCAGCT  (-7,+3)
M2.4   TGAAGAGCTGGTCTGTTGGAATAATCA...GGGTCGTTCAGCT  (-3,+0)
M2.5   TGAAGAGCTGGTCTGTTGGAATAAAAAACCGGGTCGTTCAGCT  (-0,+2)
M2.6   TGAAGAGCTGGTCTGTTGGAATAATCATATTATTATTAATCATATTTACACCGGGTCGTTCAGCT  (-0,+22)
M3.1   TGAAGAGCTGGTCTGTTGGAATAAT...CCGGGTCGTTCAGCT  (-3,+0)
M3.2   TGAAGAGCTGGTCTGTTGGAATAATCAACAAACCGGGTCGTTCAGCT  (-0,+4)
M3.3   TGAAGAGCTGGTCTGTTGGAATA...AACCGGGTCGTTCAGCT  (-3,+0)
M3.4   TGAAGAGCTGGTCTGTTGGAATAATCAATTAATAATAAACCGGGTCGTTCAGCT  (-0,+11)

p2rx5_Target_site_1
WT     GTCATTGACACAGGAGCATGGCTCAGACCTGGGGTAACTTTTTCTTCTCTCTA
M4.1   GTCATTGACACAGGAGCATGGCA.......TGGGTAACTTTTTCTTCTCTCTA  (-9,+2)
M4.2   GTCATTGACACAGGAGCATGGC......CTGGGGTAACTTTTTCTTCTCTCTA  (-6,+0)
M4.3   GTCATTGACACAGGAGCATGGCTC.....TGGGGTAACTTTTTCTTCTCTCTA  (-5,+0)
M4.4   GTCATTGACACAGGAGCATGG.............TAACTTTTTCTTCTCTCTA  (-13,+0)
M4.5   GTCATTGACACAGGAGCATGGCTC....CTGGGGTAACTTTTTCTTCTCTCTA  (-4,+0)
M4.6   GTCATTGACACTGGAGCATGGCTCAGTTACCTGGGGTAACTTTTTCTTCTCTCTA  (-1,+3)
M4.7   GTCATTGACACAGGAGCATGGCTCAGATTGACAAAGGGGTAACTTTTTCTTCTCTCTA  (-3,+8)
M5.1   GTCATTGACACAGGAGCATGGCTGTAACTTTGGGTAACTTTTTCTTCTCTCTA  (-8,+8)
M5.2   GTCATTGACACAGGAGCATGGCT.......GGGGTAACTTTTTCTTCTCTCTA  (-7,+0)
M5.3   GTCATTGACACAGGAGCCTGG...........GGTAACTTTTTCTTCTCTCTA  (-11,+0)
M5.4   GTCATTGACACAGGAGCATGGCTCA......GGGTAACTTTTTCTTCTCTCTA  (-6,+0)
M5.5   GTCATTGACACAGGAGCAT...........GGGGTAACTTTTTCTTCTCTCTA  (-11,+0)
M5.6   GTCATTGACACAGGAGCATGGCTGTGTGGCTGGGGTAACTTTTTCTTCTCTCTA  (-5,+6)
M5.7   GTCATTGACACAGGAGCATGGCTCAGGAGCAAGTTACCTGGGGTAACTTTTTCTTCTCTCTA  (-0,+9)

p2rx5_Target_site_2
WT     TTTCAGCTGACGGTGATCGGATATCTGATCGGGTAGGTGCCTT←16 bp→AGACGTTAAGGTGAACG←126 bp→CGACAGTGAACTGTTTATTA
M6.1   TTTCAGCTGACGGTGATCGGATATCT←33 bp→AGACGTTAAGGTGAACG  (-33,+0)
M6.2   TTTCAGCTGACGGTGATCGGATATCTG←175 bp→CGACAGTGAACTGTTTATTA  (-175,+0)
M6.3   TTTCAGCTGACGGTGATCGGAT.......CGGGTAGGTGCCTT  (-7,+0)
M6.4   TTTCAGCTGACGGTGATCGGGTA............GGTGCCTT  (-12,+0)
M6.5   TTTCAGCTGACGGTGATCGGA............TAGGTGCCTT  (-12,+0)
```

**Supplementary Figure 13.** Targeted indel mutations induced by CRISPR/Cas9 system of zebrafish *p2rx3a* and *p2rx5* genes.

Zebrafish *p2rx3a* and *p2rx5* were targeted. Target sequences of representative embryos were amplified, cloned and sequenced. Allele sequences with indel mutations are shown, and wild type (WT) alleles are not shown. The green letters indicate the protospacer adjacent motif (PAM) and target site in WT reference sequence, and the target sites highlighted in yellow. Deletions are shown as dashes, while large deletions are marked with the length of the missing sequences between double sided arrows. Insertions were font in red. Numbers in brackets shows the number of nucleotides deleted (−) or inserted (+) in the edited genes..

### Supplementary Tables

**Supplementary Table 1.** Summary of the sequencing data from WGS libraries and Hi-C libraries

| Strategy | Insert Size (bp) | Reads Length (bp) | Total Data (Gb) |
|---|---|---|---|
| WGS | 170 | 100_100 | 383.98 |
| | 350 | 100_100 | 387.24 |
| | 2,000 | 50_50 | 65.36 |
| | 5,000 | 50_50 | 142.20 |
| | 10,000 | 50_50 | 71.34 |
| | 20,000 | 50_50 | 57.56 |
| HIC | \ | 50_50 | 71.95 |
| Total | \ | \ | 1179.63 |

**Supplementary Table 2.** Basic statistics of the assembled bamboo shark genome

| | WGS | | HIC | |
|---|---|---|---|---|
| | Contig (bp) | Scaffold (bp) | Contig (bp) | Scaffold (bp) |
| N90 | 1,473 | 2,360 | 1,415 | 2,218 |
| N80 | 9,471 | 172,678 | 8,377 | 20,708,781 |
| N70 | 18,406 | 430,414 | 17,000 | 44,175,687 |
| N60 | 27,604 | 684,074 | 25,851 | 49,995,320 |
| N50 | 37,247 | 962,547 | 35,451 | 57,918,702 |
| N40 | 48,650 | 1,251,191 | 46,889 | 71,834,548 |
| N30 | 62,981 | 1,595,392 | 60,770 | 80,155,495 |
| N20 | 82,569 | 2,081,779 | 80,154 | 93,009,420 |
| N10 | 115,410 | 2,827,051 | 112,378 | 99,330,644 |
| max_length | 7,042,168 | 438,259 | 438,259 | 136,536,87 6 |
| Total length | 3,647,281,518 | 3,846,300,841 | 3,647,255,807 | 3,851,610,035 |

**Supplementary Table 3.** Evaluation of the final gene set by using BUSCO

| Description | Gene number | Percent (%) |
|---|---|---|
| **Complete BUSCOs (C)** | 2,477 | 95.78 |

| | | |
|---|---|---|
| **Complete and single-copy BUSCOs (S)** | 2,372 | 91.72 |
| **Complete and duplicated BUSCOs (D)** | 105 | 4.06 |
| **Fragmented BUSCOs (F)** | 68 | 2.62 |
| **Missing BUSCOs (M)** | 41 | 1.59 |
| **Total BUSCO groups searched** | 2,586 | 100.00 |

**Supplementary Table 4.** Summary of the functional annotation of genes in bamboo shark genome

| | **Number** | **Percent (%)** |
|---|---|---|
| **Total** | 19,595 | 100 |
| **InterPro** | 16,938 | 86.44 |
| **GO** | 13,147 | 67.09 |
| **KEGG** | 17,171 | 87.63 |
| **Swiss-Prot** | 18,507 | 94.45 |
| **TrEMBL** | 15,622 | 79.72 |
| **Annotated** | 19,371 | 98.86 |
| **Unannotated** | 224 | 1.24 |

**Supplementary Table 5.** Summary of the repeat content in bamboo shark genome

| **Type** | **Repeat Size** | **% of genome** |
|---|---|---|
| **Trf** | 193,033,335 | 5.01 |
| **Repeatmasker** | 945,921,870 | 24.56 |
| **Proteinmask** | 855,315,381 | 22.24 |
| **De novo** | 2,266,569,286 | 58.85 |
| **Total** | 2,447,074,981 | 63.53 |

**Supplementary Table 6.** Summary of the TE content in three cartilaginous fishes and seven bony fishes.

| | **Type** | **DNA** | **LINE** | **SINE** | **LTR** | **Total** |
|---|---|---|---|---|---|---|
| | Length (bp) | 84,246,842 | 1,841,705,606 | 151,630,200 | 707,421,972 | 2,240,930,640 |

| | | | | | | |
|---|---|---|---|---|---|---|
| Bamboo shark | % in genome | 2.19 | 47.88 | 3.94 | 18.39 | 58.26 |
| Elephant shark | Length (bp) | 20,045,841 | 375,567,981 | 150,517,703 | 119,161,620 | 444,674,297 |
| | % in genome | 2.06 | 38.54 | 15.45 | 12.23 | 45.63 |
| Whale shark | Length (bp) | 44,888,512 | 1,482,979,651 | 102,877,514 | 120,791,609 | 1,608,053,993 |
| | % in genome | 1.53 | 50.59 | 3.51 | 4.12 | 54.85 |
| Coelacanth | Length (bp) | 682,694,273 | 654,465,678 | 345,675,101 | 219,465,253 | 1,253,477,861 |
| | % in genome | 23.87 | 22.88 | 12.08 | 7.67 | 43.82 |
| Three-spined sticklebacks | Length (bp) | 35,650,096 | 28,697,415 | 3,533,002 | 30,466,795 | 82,399,819 |
| | % in genome | 7.72 | 6.22 | 0.77 | 6.6 | 17.85 |
| Large yellow croaker | Length (bp) | 50,367,111 | 26,418,884 | 3,139,446 | 17,332,054 | 85,823,340 |
| | % in genome | 7.77 | 4.07 | 0.48 | 2.67 | 13.24 |
| Medaka | Length (bp) | 126,716,317 | 90,435,719 | 9,001,570 | 62,654,967 | 253,215,905 |
| | % in genome | 14.58 | 10.41 | 1.04 | 7.21 | 29.14 |
| Torafugu | Length (bp) | 21,275,275 | 22,316,804 | 1,056,847 | 15,374,370 | 49,743,515 |
| | % in genome | 5.43 | 5.7 | 0.27 | 3.93 | 12.71 |
| Zebrafish | Length (bp) | 599,266,189 | 59,287,717 | 36,817,925 | 79,463,691 | 744,738,398 |
| | % in genome | 43.69 | 4.32 | 2.68 | 5.79 | 54.29 |
| Tongue soles | Length (bp) | 36,780,534 | 17,599,748 | 1,876,835 | 11,627,464 | 59,151,167 |
| | % in genome | 7.82 | 3.74 | 0.4 | 2.47 | 12.58 |

**Supplementary Table 7.** Distribution of paralogous genes in the bamboo shark genome. The yellow highlighted numbers represent gene pairs more than 20.

| chr | 1 | 10 | 11 | 12 | 13 | 14 | 15 | 16 | 17 | 18 | 19 | 2 | 20 | 21 | 22 | 23 | 24 | 25 | 26 | 27 | 28 | 29 | 3 | 30 | 31 | 32 | 33 | 34 | 35 | 36 | 37 | 38 | 39 | 4 | 40 | 41 | 42 | 43 | 44 | 45 | 46 | 47 | 48 | 49 | 5 | 50 | 51 | 6 | 7 | 8 | 9 |
|---|---|---|---|---|---|---|---|---|---|---|---|---|---|---|---|---|---|---|---|---|---|---|---|---|---|---|---|---|---|---|---|---|---|---|---|---|---|---|---|---|---|---|---|---|---|---|---|---|---|---|---|
| 1 | 7 | 0 | 10 | 3 | 0 | 39 | 7 | 0 | 0 | 0 | 0 | 2 | 1 | 1 | 18 | 0 | 1 | 0 | 1 | 0 | 1 | 0 | 1 | 0 | 12 | 0 | 1 | 0 | 0 | 12 | 0 | 11 | 0 | 1 | 10 | 0 | 7 | 0 | 0 | 0 | 0 | 0 | 0 | 0 | 1 | 0 | 2 | 25 | 0 | 0 | 0 |
| 10 | | 10 | 0 | 0 | 1 | 1 | 0 | 0 | 0 | 1 | 1 | 0 | 0 | 0 | 0 | 0 | 0 | 0 | 0 | 0 | 0 | 0 | 18 | 0 | 0 | 1 | 0 | 0 | 0 | 0 | 0 | 0 | 1 | 1 | 0 | 25 | 0 | 0 | 1 | 6 | 0 | 1 | 0 | 1 | 2 | 0 | 0 | 0 | 0 | 0 | 58 |
| 11 | | | 7 | 0 | 0 | 0 | 0 | 0 | 0 | 0 | 0 | 17 | 0 | 1 | 0 | 0 | 0 | 1 | 0 | 0 | 0 | 1 | 2 | 36 | 7 | 1 | 0 | 5 | 0 | 0 | 2 | 0 | 0 | 0 | 0 | 0 | 0 | 0 | 0 | 0 | 0 | 0 | 0 | 0 | 1 | 0 | 0 | 0 | 2 | 27 | 0 |
| 12 | | | | 9 | 7 | 0 | 21 | 0 | 0 | 2 | 0 | 2 | 0 | 0 | 2 | 0 | 2 | 0 | 0 | 6 | 0 | 2 | 8 | 1 | 0 | 0 | 1 | 0 | 18 | 1 | 0 | 0 | 0 | 0 | 0 | 0 | 0 | 0 | 0 | 0 | 0 | 0 | 0 | 0 | 0 | 0 | 0 | 0 | 0 | 0 | 8 |
| 13 | | | | | 19 | 0 | 11 | 0 | 1 | 0 | 0 | 0 | 0 | 0 | 0 | 0 | 0 | 1 | 1 | 0 | 0 | 0 | 0 | 0 | 0 | 0 | 0 | 28 | 0 | 0 | 0 | 0 | 0 | 0 | 0 | 0 | 0 | 2 | 0 | 0 | 0 | 0 | 13 | 0 | 0 | 0 | 0 | 21 | 3 | 1 | 0 |
| 14 | | | | | | 8 | 3 | 0 | 0 | 1 | 0 | 13 | 0 | 0 | 10 | 0 | 0 | 0 | 0 | 1 | 0 | 0 | 0 | 1 | 0 | 26 | 0 | 0 | 0 | 0 | 0 | 8 | 0 | 0 | 0 | 1 | 5 | 0 | 0 | 0 | 0 | 0 | 0 | 0 | 0 | 0 | 1 | 9 | 0 | 0 | 0 |
| 15 | | | | | | | 3 | 0 | 2 | 0 | 0 | 3 | 1 | 0 | 1 | 0 | 0 | 0 | 0 | 2 | 1 | 0 | 2 | 1 | 1 | 4 | 0 | 0 | 0 | 0 | 0 | 0 | 1 | 0 | 0 | 0 | 0 | 0 | 0 | 0 | 0 | 0 | 2 | 0 | 0 | 0 | 0 | 60 | 0 | 0 | 0 |
| 16 | | | | | | | | 5 | 0 | 0 | 26 | 0 | 0 | 0 | 0 | 12 | 0 | 1 | 10 | 0 | 0 | 0 | 0 | 0 | 0 | 0 | 0 | 0 | 0 | 0 | 0 | 0 | 17 | 1 | 2 | 1 | 0 | 0 | 0 | 0 | 0 | 0 | 0 | 0 | 2 | 0 | 0 | 0 | 0 | 0 | 0 |
| 17 | | | | | | | | | 10 | 0 | 0 | 1 | 7 | 0 | 0 | 0 | 0 | 1 | 0 | 1 | 0 | 13 | 1 | 0 | 0 | 0 | 0 | 0 | 0 | 1 | 0 | 0 | 0 | 24 | 0 | 0 | 0 | 1 | 0 | 0 | 0 | 0 | 0 | 0 | 7 | 0 | 0 | 0 | 1 | 0 | 0 |
| 18 | | | | | | | | | | 7 | 39 | 1 | 1 | 0 | 0 | 25 | 0 | 0 | 11 | 0 | 1 | 0 | 1 | 0 | 0 | 0 | 0 | 0 | 0 | 0 | 0 | 0 | 1 | 0 | 0 | 0 | 0 | 0 | 0 | 0 | 0 | 0 | 0 | 0 | 0 | 0 | 0 | 0 | 0 | 0 | 0 |
| 19 | | | | | | | | | | | 3 | 0 | 0 | 0 | 0 | 0 | 0 | 0 | 41 | 0 | 1 | 0 | 0 | 1 | 0 | 1 | 1 | 0 | 2 | 0 | 0 | 0 | 5 | 0 | 0 | 1 | 1 | 0 | 0 | 0 | 0 | 0 | 0 | 0 | 0 | 0 | 0 | 0 | 0 | 0 | 0 |
| 2 | | | | | | | | | | | | 13 | 0 | 0 | 6 | 0 | 0 | 0 | 0 | 0 | 0 | 0 | 0 | 0 | 53 | 8 | 0 | 0 | 0 | 19 | 0 | 0 | 0 | 0 | 18 | 0 | 3 | 0 | 0 | 0 | 0 | 0 | 0 | 0 | 0 | 0 | 4 | 1 | 1 | 2 | 0 |
| 20 | | | | | | | | | | | | | 5 | 0 | 0 | 0 | 0 | 0 | 0 | 3 | 0 | 5 | 8 | 0 | 0 | 0 | 0 | 0 | 0 | 0 | 0 | 0 | 0 | 66 | 0 | 0 | 0 | 0 | 0 | 0 | 0 | 0 | 0 | 0 | 5 | 0 | 0 | 0 | 0 | 0 | 2 |
| 21 | | | | | | | | | | | | | | 8 | 0 | 0 | 45 | 2 | 0 | 0 | 1 | 0 | 0 | 0 | 0 | 0 | 0 | 0 | 0 | 0 | 0 | 0 | 20 | 0 | 0 | 0 | 0 | 0 | 0 | 0 | 0 | 0 | 0 | 0 | 0 | 0 | 0 | 0 | 3 | 0 | 0 |
| 22 | | | | | | | | | | | | | | | 4 | 1 | 0 | 0 | 0 | 0 | 0 | 0 | 0 | 0 | 0 | 8 | 0 | 0 | 0 | 0 | 0 | 0 | 0 | 1 | 0 | 0 | 25 | 0 | 0 | 0 | 0 | 0 | 0 | 0 | 0 | 0 | 0 | 0 | 0 | 0 | 0 |
| 23 | | | | | | | | | | | | | | | | 7 | 0 | 0 | 31 | 1 | 1 | 0 | 0 | 0 | 0 | 0 | 1 | 0 | 0 | 0 | 0 | 0 | 5 | 0 | 0 | 0 | 0 | 0 | 0 | 0 | 0 | 0 | 0 | 0 | 0 | 0 | 0 | 0 | 0 | 1 | 1 |
| 24 | | | | | | | | | | | | | | | | | 6 | 0 | 0 | 0 | 0 | 0 | 0 | 0 | 0 | 0 | 0 | 0 | 0 | 0 | 0 | 0 | 11 | 0 | 0 | 0 | 0 | 0 | 0 | 1 | 0 | 0 | 0 | 0 | 1 | 0 | 0 | 1 | 0 | 0 | 0 |
| 25 | | | | | | | | | | | | | | | | | | 10 | 0 | 1 | 52 | 1 | 1 | 0 | 0 | 0 | 0 | 1 | 0 | 0 | 0 | 0 | 0 | 0 | 0 | 1 | 0 | 0 | 1 | 0 | 1 | 0 | 0 | 0 | 0 | 0 | 0 | 0 | 0 | 0 | 0 |
| 26 | | | | | | | | | | | | | | | | | | | 10 | 1 | 0 | 0 | 1 | 0 | 0 | 0 | 0 | 0 | 0 | 0 | 0 | 0 | 4 | 0 | 0 | 0 | 0 | 0 | 0 | 0 | 0 | 0 | 0 | 0 | 0 | 0 | 0 | 0 | 0 | 0 | 0 |
| 27 | | | | | | | | | | | | | | | | | | | | 7 | 0 | 1 | 42 | 0 | 0 | 0 | 1 | 0 | 0 | 0 | 0 | 0 | 0 | 7 | 0 | 0 | 0 | 0 | 0 | 0 | 0 | 0 | 0 | 0 | 0 | 0 | 0 | 0 | 0 | 0 | 12 |
| 28 | | | | | | | | | | | | | | | | | | | | | 5 | 1 | 0 | 0 | 0 | 0 | 0 | 1 | 0 | 0 | 0 | 0 | 2 | 0 | 0 | 0 | 0 | 0 | 0 | 0 | 0 | 0 | 0 | 0 | 0 | 0 | 0 | 1 | 1 | 0 | 0 |
| 29 | | | | | | | | | | | | | | | | | | | | | | 8 | 5 | 0 | 0 | 0 | 2 | 0 | 2 | 0 | 0 | 0 | 0 | 1 | 0 | 0 | 0 | 0 | 0 | 0 | 0 | 0 | 0 | 0 | 1 | 0 | 0 | 1 | 4 | 0 | 1 |
| 3 | | | | | | | | | | | | | | | | | | | | | | | 17 | 0 | 0 | 0 | 0 | 0 | 1 | 0 | 1 | 1 | 0 | 25 | 0 | 2 | 0 | 0 | 0 | 7 | 0 | 0 | 0 | 0 | 0 | 0 | 0 | 9 | 1 | 0 | 0 |
| 30 | | | | | | | | | | | | | | | | | | | | | | | | 2 | 0 | 0 | 0 | 2 | 0 | 0 | 5 | 0 | 0 | 1 | 0 | 0 | 0 | 0 | 0 | 0 | 0 | 0 | 0 | 0 | 0 | 0 | 0 | 0 | 0 | 9 | 0 |
| 31 | | | | | | | | | | | | | | | | | | | | | | | | | 11 | 2 | 0 | 0 | 0 | 5 | 0 | 0 | 0 | 0 | 10 | 0 | 0 | 0 | 0 | 0 | 0 | 0 | 0 | 0 | 0 | 0 | 0 | 1 | 0 | 4 | 0 |
| 32 | | | | | | | | | | | | | | | | | | | | | | | | | | 2 | 0 | 0 | 0 | 0 | 0 | 3 | 0 | 0 | 0 | 0 | 3 | 0 | 0 | 0 | 0 | 0 | 0 | 0 | 1 | 0 | 1 | 3 | 0 | 0 | 0 |
| 33 | | | | | | | | | | | | | | | | | | | | | | | | | | | 4 | 1 | 0 | 0 | 0 | 0 | 0 | 9 | 0 | 0 | 0 | 5 | 0 | 0 | 0 | 0 | 0 | 0 | 42 | 0 | 0 | 0 | 13 | 0 | 0 |
| 34 | | | | | | | | | | | | | | | | | | | | | | | | | | | | 2 | 0 | 0 | 0 | 0 | 0 | 0 | 0 | 0 | 0 | 1 | 0 | 0 | 0 | 0 | 0 | 0 | 0 | 0 | 0 | 0 | 0 | 0 | 0 |
| 35 | | | | | | | | | | | | | | | | | | | | | | | | | | | | | 1 | 0 | 1 | 0 | 0 | 1 | 0 | 0 | 0 | 0 | 0 | 0 | 1 | 0 | 0 | 0 | 0 | 0 | 0 | 3 | 0 | 0 | 1 |
| 36 | | | | | | | | | | | | | | | | | | | | | | | | | | | | | | 2 | 0 | 0 | 1 | 0 | 0 | 0 | 0 | 0 | 0 | 0 | 0 | 0 | 0 | 0 | 0 | 0 | 3 | 0 | 0 | 0 | 0 |
| 37 | | | | | | | | | | | | | | | | | | | | | | | | | | | | | | | 20 | 0 | 0 | 0 | 0 | 0 | 0 | 0 | 0 | 0 | 1 | 0 | 0 | 0 | 1 | 0 | 0 | 0 | 0 | 1 | 0 |
| 38 | | | | | | | | | | | | | | | | | | | | | | | | | | | | | | | | 3 | 0 | 1 | 1 | 0 | 17 | 0 | 0 | 0 | 0 | 0 | 0 | 0 | 0 | 1 | 0 | 0 | 0 | 0 | 0 |
| 39 | | | | | | | | | | | | | | | | | | | | | | | | | | | | | | | | | 3 | 0 | 0 | 0 | 0 | 0 | 0 | 0 | 0 | 0 | 0 | 0 | 0 | 0 | 0 | 1 | 0 | 0 | 0 |
| 4 | | | | | | | | | | | | | | | | | | | | | | | | | | | | | | | | | | 4 | 0 | 0 | 0 | 6 | 0 | 0 | 0 | 1 | 0 | 0 | 0 | 0 | 0 | 0 | 11 | 1 | 3 |
| 40 | | | | | | | | | | | | | | | | | | | | | | | | | | | | | | | | | | | 5 | 0 | 0 | 0 | 0 | 0 | 0 | 0 | 0 | 0 | 0 | 0 | 2 | 1 | 0 | 0 | 0 |
| 41 | | | | | | | | | | | | | | | | | | | | | | | | | | | | | | | | | | | | 3 | 0 | 0 | 0 | 1 | 0 | 0 | 0 | 0 | 0 | 0 | 0 | 0 | 0 | 0 | 9 |
| 42 | | | | | | | | | | | | | | | | | | | | | | | | | | | | | | | | | | | | | 2 | 0 | 0 | 0 | 3 | 0 | 1 | 0 | 0 | 0 | 0 | 0 | 2 | 0 | 0 |
| 43 | | | | | | | | | | | | | | | | | | | | | | | | | | | | | | | | | | | | | | 1 | 0 | 0 | 0 | 0 | 0 | 0 | 6 | 0 | 0 | 0 | 22 | 0 | 0 |
| 44 | | | | | | | | | | | | | | | | | | | | | | | | | | | | | | | | | | | | | | | 18 | 3 | 1 | 0 | 0 | 0 | 0 | 0 | 0 | 0 | 0 | 0 | 0 |
| 45 | | | | | | | | | | | | | | | | | | | | | | | | | | | | | | | | | | | | | | | | 4 | 0 | 0 | 0 | 0 | 0 | 0 | 0 | 0 | 0 | 0 | 10 |
| 46 | | | | | | | | | | | | | | | | | | | | | | | | | | | | | | | | | | | | | | | | | 4 | 0 | 0 | 0 | 0 | 0 | 0 | 0 | 0 | 2 | 0 |
| 47 | | | | | | | | | | | | | | | | | | | | | | | | | | | | | | | | | | | | | | | | | | 3 | 0 | 0 | 0 | 0 | 0 | 0 | 0 | 0 | 0 |
| 48 | | | | | | | | | | | | | | | | | | | | | | | | | | | | | | | | | | | | | | | | | | | 3 | 0 | 0 | 0 | 0 | 1 | 1 | 0 | 0 |
| 49 | | | | | | | | | | | | | | | | | | | | | | | | | | | | | | | | | | | | | | | | | | | | 0 | 0 | 0 | 0 | 0 | 0 | 0 | 0 |
| 5 | | | | | | | | | | | | | | | | | | | | | | | | | | | | | | | | | | | | | | | | | | | | | 8 | 0 | 0 | 0 | 59 | 0 | 0 |
| 50 | | | | | | | | | | | | | | | | | | | | | | | | | | | | | | | | | | | | | | | | | | | | | | 6 | 0 | 0 | 0 | 0 | 0 |
| 51 | | | | | | | | | | | | | | | | | | | | | | | | | | | | | | | | | | | | | | | | | | | | | | | 0 | 0 | 0 | 0 | 2 |
| 6 | | | | | | | | | | | | | | | | | | | | | | | | | | | | | | | | | | | | | | | | | | | | | | | | 13 | 0 | 0 | 1 |
| 7 | | | | | | | | | | | | | | | | | | | | | | | | | | | | | | | | | | | | | | | | | | | | | | | | | 9 | 0 | 0 |
| 8 | | | | | | | | | | | | | | | | | | | | | | | | | | | | | | | | | | | | | | | | | | | | | | | | | | 7 | 0 |
| 9 | | | | | | | | | | | | | | | | | | | | | | | | | | | | | | | | | | | | | | | | | | | | | | | | | | | 12 |

**Supplementary Table 8.** Summary of conserved genes on each chromosome.

| Chr | Total gene number | Conserved gene number | Conserved gene length (bp) | Chr length (bp) | Percent of conserved gene length (%) | BUSCO gene number | Percent of BUSCO genes (%) |
|---|---|---|---|---|---|---|---|
| 1 | 749 | 398 | 26,983,328 | 156,605,781 | 17.23 | 115 | 0.15 |
| 2 | 671 | 362 | 27,059,668 | 146,811,608 | 18.43 | 119 | 0.18 |
| 3 | 714 | 418 | 29,379,912 | 138,146,672 | 21.27 | 135 | 0.19 |
| 4 | 650 | 359 | 28,469,618 | 138,123,871 | 20.61 | 115 | 0.18 |

| | | | | | | | |
|---|---|---|---|---|---|---|---|
| 5 | 575 | 310 | 26,195,248 | 124,146,585 | 21.10 | 83 | 0.14 |
| 6 | 585 | 338 | 23,777,413 | 121,738,290 | 19.53 | 102 | 0.17 |
| 7 | 622 | 345 | 22,949,759 | 118,101,210 | 19.43 | 92 | 0.15 |
| 8 | 243 | 59 | 2,374,595 | 107,642,081 | 2.21 | 6 | 0.02 |
| 9 | 558 | 335 | 26,847,317 | 105,880,166 | 25.36 | 105 | 0.19 |
| 10 | 538 | 327 | 21,741,870 | 95,765,976 | 22.70 | 106 | 0.20 |
| 11 | 572 | 329 | 20,875,770 | 94,731,789 | 22.04 | 99 | 0.17 |
| 12 | 518 | 287 | 17,276,967 | 88,215,905 | 19.58 | 116 | 0.22 |
| 13 | 528 | 277 | 15,199,283 | 84,432,200 | 18.00 | 73 | 0.14 |
| 14 | 412 | 217 | 10,918,334 | 82,264,610 | 13.27 | 49 | 0.12 |
| 15 | 397 | 210 | 10,794,017 | 81,258,972 | 13.28 | 34 | 0.09 |
| 16 | 394 | 179 | 13,608,314 | 76,207,864 | 17.86 | 56 | 0.14 |
| 17 | 441 | 275 | 16,695,914 | 73,167,756 | 22.82 | 76 | 0.17 |
| 18 | 404 | 206 | 13,036,022 | 72,140,527 | 18.07 | 63 | 0.16 |
| 19 | 430 | 263 | 15,698,935 | 69,975,007 | 22.44 | 91 | 0.21 |
| 20 | 362 | 203 | 10,412,038 | 64,470,864 | 16.15 | 65 | 0.18 |
| 21 | 446 | 264 | 14,805,954 | 60,475,494 | 24.48 | 91 | 0.20 |
| 22 | 366 | 229 | 14,577,723 | 59,434,692 | 24.53 | 78 | 0.21 |
| 23 | 350 | 216 | 14,198,713 | 58,390,607 | 24.32 | 65 | 0.19 |
| 24 | 357 | 189 | 10,959,335 | 56,061,091 | 19.55 | 49 | 0.14 |
| 25 | 386 | 238 | 13,505,391 | 54,600,476 | 24.73 | 61 | 0.16 |
| 26 | 325 | 147 | 8,503,535 | 53,348,395 | 15.94 | 32 | 0.10 |
| 27 | 303 | 179 | 7,205,788 | 51,668,882 | 13.95 | 44 | 0.15 |
| 28 | 348 | 203 | 13,925,264 | 51,251,456 | 27.17 | 68 | 0.20 |
| 29 | 249 | 150 | 9,496,342 | 49,130,325 | 19.33 | 53 | 0.21 |
| 30 | 324 | 171 | 9,589,896 | 47,581,304 | 20.15 | 48 | 0.15 |
| 31 | 389 | 203 | 9,051,487 | 45,411,694 | 19.93 | 23 | 0.06 |
| 32 | 255 | 134 | 7,083,153 | 45,129,243 | 15.70 | 49 | 0.19 |
| 33 | 307 | 149 | 7,312,983 | 43,998,411 | 16.62 | 23 | 0.07 |
| 34 | 311 | 180 | 10,508,857 | 43,401,492 | 24.21 | 54 | 0.17 |
| 35 | 217 | 93 | 5,442,497 | 42,304,929 | 12.86 | 29 | 0.13 |
| 36 | 239 | 132 | 6,918,362 | 38,655,811 | 17.90 | 40 | 0.17 |
| 37 | 360 | 25 | 1,803,637 | 35,007,441 | 5.15 | 7 | 0.02 |

| | | | | | | | |
|---|---|---|---|---|---|---|---|
| 38 | 205 | 118 | 6,059,768 | 30,225,729 | 20.05 | 21 | 0.10 |
| 39 | 340 | 95 | 2,836,048 | 27,915,349 | 10.16 | 21 | 0.06 |
| 40 | 205 | 122 | 7,002,794 | 26,992,261 | 25.94 | 28 | 0.14 |
| 41 | 212 | 54 | 2,134,626 | 20,367,195 | 10.48 | 5 | 0.02 |
| 42 | 225 | 88 | 2,767,007 | 19,796,787 | 13.98 | 19 | 0.08 |
| 43 | 149 | 34 | 1,360,873 | 16,392,901 | 8.30 | 5 | 0.03 |
| 44 | 177 | 0 | 0 | 15,333,346 | 0.00 | 0 | 0.00 |
| 45 | 188 | 12 | 214,271 | 13,660,452 | 1.57 | 1 | 0.01 |
| 46 | 57 | 7 | 447,174 | 12,823,346 | 3.49 | 1 | 0.02 |
| 47 | 73 | 3 | 14,540 | 10,712,325 | 0.14 | 0 | 0.00 |
| 48 | 124 | 18 | 345,936 | 9,987,384 | 3.46 | 2 | 0.02 |
| 49 | 13 | 1 | 7,795 | 9,983,820 | 0.08 | 0 | 0.00 |
| 50 | 49 | 0 | 0 | 8,079,144 | 0.00 | 0 | 0.00 |
| 51 | 97 | 12 | 203,648 | 4,637,688 | 4.39 | 7 | 0.07 |

**Supplementary Table 9.** Enrichment analysis for genes in chromosomes 8, 37, 39, 41, 43, 44, 45, 46, 47,48,49, 50 and 51.

| **Pathway** | **Level** | **Gene number** | **Q-value** |
|---|---|---|---|
| Systemic lupus erythematosus | Immune diseases | 66 | 6.08E-23 |
| Staphylococcus aureus infection | Infectious diseases: Bacterial | 47 | 1.40E-14 |
| Asthma | Immune diseases | 36 | 1.84E-14 |
| Intestinal immune network for IgA production | Immune system | 40 | 4.66E-14 |
| Malaria | Infectious diseases: Parasitic | 37 | 1.05E-12 |
| Measles | Infectious diseases: Viral | 59 | 2.02E-10 |
| Mineral absorption | Digestive system | 48 | 3.52E-10 |
| Allograft rejection | Immune diseases | 36 | 1.15E-09 |

| | | | |
|---|---|---|---|
| NF-kappa B signaling pathway | Signal transduction | 54 | 1.34E-09 |
| Autoimmune thyroid disease | Immune diseases | 37 | 6.19E-09 |
| Graft-versus-host disease | Immune diseases | 28 | 1.25E-08 |
| Inflammatory bowel disease (IBD) | Immune diseases | 32 | 4.87E-08 |
| T cell receptor signaling pathway | Immune system | 45 | 6.48E-08 |
| Chagas disease (American trypanosomiasis) | Infectious diseases: Parasitic | 43 | 7.64E-08 |
| Type I diabetes mellitus | Endocrine and metabolic diseases | 28 | 2.04E-07 |
| Viral myocarditis | Cardiovascular diseases | 40 | 2.04E-06 |
| Rheumatoid arthritis | Immune diseases | 41 | 2.39E-06 |
| Antigen processing and presentation | Immune system | 34 | 6.26E-06 |
| Cell adhesion molecules (CAMs) | Signaling molecules and interaction | 51 | 5.33E-05 |
| Alcoholism | Substance dependence | 39 | 5.71E-05 |
| HTLV-I infection | Infectious diseases: Viral | 63 | 2.90E-04 |
| Transcriptional misregulation in cancer | Cancers: Overview | 56 | 7.29E-04 |
| Ras signaling pathway | Signal transduction | 56 | 7.22E-03 |
| Calcium signaling pathway | Signal transduction | 52 | 8.39E-03 |
| Leishmaniasis | Infectious diseases: Parasitic | 22 | 8.68E-03 |
| Rap1 signaling pathway | Signal transduction | 57 | 8.90E-03 |
| Viral carcinogenesis | Cancers: Overview | 44 | 2.35E-02 |
| Tuberculosis | Infectious diseases: Bacterial | 37 | 6.91E-02 |

**Supplementary Table 10.** List of immune-related genes in chromosomes 8, 37, 39, 41, 43, 44, 45, 46, 47, 48, 49, 50 and 51.

| Chromosomes | Gene ID | Descriptions from KEGG database |
|---|---|---|
| | Chipl18455 | Ig mu chain C region secreted form; K06856 immunoglobulin heavy chain |
| | Chipl18459 | cathepsin L1-like; K01365 cathepsin L [EC:3.4.22.15] |
| | Chipl18607 | AAEL011167-PA; K01365 cathepsin L [EC:3.4.22.15] |
| | Chipl18610 | uncharacterized LOC548369; K06856 immunoglobulin heavy chain |
| | Chipl18619 | IgX; Ig mu chain C region membrane-bound form; K06856 immunoglobulin heavy chain |
| | Chipl18628 | T-cell receptor beta-1 chain C region; K10785 T-cell receptor beta chain V region |
| | Chipl18778 | MAP2K7; mitogen-activated protein kinase kinase 7; K04431 mitogen-activated protein kinase kinase 7 [EC:2.7.12.2] |
| | Chipl18792 | cAMP-dependent protein kinase catalytic subunit alpha; K04345 protein kinase A [EC:2.7.11.11] |
| | Chipl18812 | hypothetical protein; K08059 interferon, gamma-inducible protein 30 |
| | Chipl18816 | DNA-directed RNA polymerase III subunit RPC2-like; K03021 DNA-directed RNA polymerase III subunit RPC2 [EC:2.7.7.6] |
| | Chipl18831 | lipid phosphate phosphohydrolase 3-like; K01080 phosphatidate phosphatase [EC:3.1.3.4] |
| | Chipl18853 | EPOR; erythropoietin receptor; K05079 erythropoietin receptor |
| | Chipl18864 | calr; calreticulin; K08057 calreticulin |
| | Chipl18895 | c3; complement C3; K03990 complement component 3 |
| | Chipl18896 | c3; complement C3; K03990 complement component 3 |
| | Chipl18897 | c3; complement C3; K03990 complement component 3 |
| chr8 | Chipl18903 | pin1; peptidylprolyl cis/trans isomerase, NIMA-interacting 1; K09578 peptidyl-prolyl cis-trans isomerase NIMA-interacting 1 [EC:5.2.1.8] |

| | | |
|---|---|---|
| | Chipl18905 | VAV1; vav guanine nucleotide exchange factor 1; K05730 guanine nucleotide exchange factor VAV |
| | Chipl18945 | vascular cell adhesion protein 1-like; K06527 vascular cell adhesion molecule 1 |
| | Chipl18946 | vascular cell adhesion protein 1-like; K06527 vascular cell adhesion molecule 1 |
| | Chipl18947 | vascular cell adhesion protein 1-like; K06527 vascular cell adhesion molecule 1 |
| | Chipl18948 | vascular cell adhesion protein 1-like; K06527 vascular cell adhesion molecule 1 |
| | Chipl18949 | vascular cell adhesion protein 1-like; K06527 vascular cell adhesion molecule 1 |
| | Chipl18958 | GNG7; G protein subunit gamma 7; K04543 guanine nucleotide-binding protein G(I)/G(S)/G(O) subunit gamma-7 |
| | Chipl12832 | Lach-UA-02; uncharacterized LOC102360671; K06751 major histocompatibility complex, class I |
| | Chipl12833 | Sasa-UBA, UBA; class I histocompatibility antigen, F10 alpha chain-like; K06751 major histocompatibility complex, class I |
| | Chipl12836 | beta-2-microglobulin-like; K08055 beta-2-microglobulin |
| | Chipl12839 | DLA class II histocompatibility antigen, DR-1 beta chain-like; K06752 major histocompatibility complex, class II |
| | Chipl12840 | RLA class II histocompatibility antigen, DP alpha-1 chain-like; K06752 major histocompatibility complex, class II |
| | Chipl12871 | CARD6; caspase recruitment domain family member 6; K12797 caspase recruitment domain-containing protein 6 |
| | Chipl12873 | Lach-UA-01; class I histocompatibility antigen, F10 alpha chain-like; K06751 major histocompatibility complex, class I |
| chr37 | Chipl12885 | TAP2; transporter 2, ATP binding cassette subfamily B member; K05654 ATP-binding cassette, subfamily B (MDR/TAP), member 3 |

| | |
|---|---|
| Chipl12886 | TAP2; transporter 2, ATP binding cassette subfamily B member; K05654 ATP-binding cassette, subfamily B (MDR/TAP), member 3 |
| Chipl12889 | complement C4-like; K03989 complement component 4 |
| Chipl12890 | complement C4-like; K03989 complement component 4 |
| Chipl12891 | complement C4-like; K03989 complement component 4 |
| Chipl12893 | complement C4-like; K03989 complement component 4 |
| Chipl12895 | class I histocompatibility antigen, F10 alpha chain; K06751 major histocompatibility complex, class I |
| Chipl12897 | class I histocompatibility antigen, F10 alpha chain; K06751 major histocompatibility complex, class I |
| Chipl12898 | Lach-UA-01; class I histocompatibility antigen, F10 alpha chain-like; K06751 major histocompatibility complex, class I |
| Chipl12906 | Lach-UA-02; uncharacterized LOC102360671; K06751 major histocompatibility complex, class I |
| Chipl12907 | RLA class II histocompatibility antigen, DP alpha-1 chain-like; K06752 major histocompatibility complex, class II |
| Chipl12912 | Lach-UA-01; class I histocompatibility antigen, F10 alpha chain-like; K06751 major histocompatibility complex, class I |
| Chipl12931 | class I histocompatibility antigen, F10 alpha chain; K06751 major histocompatibility complex, class I |
| Chipl12940 | class I histocompatibility antigen, F10 alpha chain; K06751 major histocompatibility complex, class I |
| Chipl12942 | class I histocompatibility antigen, F10 alpha chain; K06751 major histocompatibility complex, class I |
| Chipl12948 | natural cytotoxicity triggering receptor 3-like; K06743 natural cytotoxicity triggering receptor 3 |
| Chipl12949 | complement C4-like; K03989 complement component 4 |
| Chipl12958 | complement C4-like; K03989 complement component 4 |
| Chipl12973 | CARD6; caspase recruitment domain family member 6; K12797 caspase recruitment domain-containing protein 6 |
| Chipl12996 | CD244; CD244 molecule; K06582 natural killer cell receptor 2B4 |

| | | |
|---|---|---|
| | Chipl13006 | DNA-directed RNA polymerase III subunit RPC2-like; K03021 DNA-directed RNA polymerase III subunit RPC2 [EC:2.7.7.6] |
| | Chipl13036 | TNF receptor-associated factor 2-like; K03173 TNF receptor-associated factor 2 [EC:2.3.2.27] |
| | Chipl13046 | natural cytotoxicity triggering receptor 3-like; K06743 natural cytotoxicity triggering receptor 3 |
| | Chipl13055 | C2; complement C2; K01332 complement component 2 [EC:3.4.21.43] |
| | Chipl13056 | CFB; complement factor B; K01335 component factor B [EC:3.4.21.47] |
| | Chipl13135 | TNF; tumor necrosis factor; K03156 tumor necrosis factor superfamily, member 2 |
| | Chipl13136 | TNF; tumor necrosis factor; K03156 tumor necrosis factor superfamily, member 2 |
| | Chipl13140 | DNA-directed RNA polymerase III subunit RPC2-like; K03021 DNA-directed RNA polymerase III subunit RPC2 [EC:2.7.7.6] |
| | Chipl13157 | DNA-directed RNA polymerase III subunit RPC2-like; K03021 DNA-directed RNA polymerase III subunit RPC2 [EC:2.7.7.6] |
| | Chipl13164 | CLDN3; claudin 3; K06087 claudin |
| | Chipl13173 | heat shock 70 kDa protein; K03283 heat shock 70kDa protein 1/8 |
| | Chipl13175 | heat shock 70 kDa protein-like; K03283 heat shock 70kDa protein 1/8 |
| | Chipl13176 | heat shock 70 kDa protein; K03283 heat shock 70kDa protein 1/8 |
| | Chipl13177 | Hsp70; heat shock cognate 71 kDa protein-like; K03283 heat shock 70kDa protein 1/8 |
| | Chipl13182 | TRAF2; TNF receptor associated factor 2; K03173 TNF receptor-associated factor 2 [EC:2.3.2.27] |
| | Chipl02885 | cd37; CD37 molecule; K06475 CD37 antigen |
| chr39 | Chipl02922 | GRB2-related adapter protein-like; K07366 GRB2-related adaptor protein 2 |

| | |
|---|---|
| Chipl02924 | TAB1; TGF-beta activated kinase 1 (MAP3K7) binding protein 1; K04403 TAK1-binding protein 1 |
| Chipl02929 | POLR2F; RNA polymerase II subunit F; K03014 DNA-directed RNA polymerases I, II, and III subunit RPABC2 |
| Chipl02943 | interferon regulatory factor 3-like; K05411 interferon regulatory factor 3 |
| Chipl02944 | interferon regulatory factor 3-like; K05411 interferon regulatory factor 3 |
| Chipl02950 | DNA-directed RNA polymerases I, II, and III subunit RPABC5-like; K03007 DNA-directed RNA polymerases I, II, and III subunit RPABC5 |
| Chipl03014 | flt3lg; fms related tyrosine kinase 3 ligand; K05454 fms-related tyrosine kinase 3 ligand |
| Chipl03037 | F9; coagulation factor IX; K01321 coagulation factor IX (Christmas factor) [EC:3.4.21.22] |
| Chipl03070 | RAC1; ras-related C3 botulinum toxin substrate 1 (rho family, small GTP binding protein Rac1); K04392 Ras-related C3 botulinum toxin substrate 1 |
| Chipl03071 | rac1, Xrac, mig5, p21-rac1, rac, tc-25; ras-related C3 botulinum toxin substrate 1 (rho family, small GTP binding protein Rac1); K04392 Ras-related C3 botulinum toxin substrate 1 |
| Chipl03074 | NCF4; neutrophil cytosolic factor 4; K08012 neutrophil cytosolic factor 4 |
| Chipl03077 | pla2g6; phospholipase A2 group VI; K16343 calcium-independent phospholipase A2 [EC:3.1.1.4] |
| Chipl03087 | CARD11; caspase recruitment domain family member 11; K07367 caspase recruitment domain-containing protein 11 |
| Chipl03099 | hypothetical protein ; K17834 cyclic GMP-AMP synthase [EC:2.7.7.86] |
| Chipl03105 | tnfsf13b; tumor necrosis factor superfamily member 13b; K05476 tumor necrosis factor ligand superfamily member 13B |
| Chipl03107 | ICOSLG; inducible T-cell costimulator ligand; K06710 inducible T-cell co-stimulator ligand |

| | | |
|---|---|---|
| | Chipl03116 | POLR3H; RNA polymerase III subunit H; K03022 DNA-directed RNA polymerase III subunit RPC8 |
| | Chipl03173 | tab1; TGF-beta activated kinase 1 (MAP3K7) binding protein 1; K04403 TAK1-binding protein 1 |
| | Chipl03200 | interleukin-1 receptor-associated kinase 1-like; K04730 interleukin-1 receptor-associated kinase 1 [EC:2.7.11.1] |
| | Chipl03239 | uncharacterized LOC101922302; K10785 T-cell receptor beta chain V region |
| | Chipl03344 | DNA-directed RNA polymerase III subunit RPC2-like; K03021 DNA-directed RNA polymerase III subunit RPC2 [EC:2.7.7.6] |
| | Chipl03362 | PAK4; p21 (RAC1) activated kinase 4; K05734 p21-activated kinase 4 [EC:2.7.11.1] |
| | Chipl03363 | alpha-actinin-1-like; K05699 actinin alpha 1/4 |
| | Chipl03396 | nfkbib; NFKB inhibitor beta; K02581 NF-kappa-B inhibitor beta |
| | Chipl03397 | NFKBIA; NFKB inhibitor alpha; K04734 NF-kappa-B inhibitor alpha |
| | Chipl03417 | TNF receptor-associated factor 3-like; K03174 TNF receptor-associated factor 3 |
| | Chipl03418 | traf3; TNF receptor associated factor 3; K03174 TNF receptor-associated factor 3 |
| | Chipl03422 | vasodilator-stimulated phosphoprotein-like; K06274 vasodilator-stimulated phosphoprotein |
| | Chipl03439 | arhgap35; Rho GTPase activating protein 35; K05732 glucocorticoid receptor DNA-binding factor 1 |
| | Chipl03450 | ptgir; prostaglandin I2 (prostacyclin) receptor (IP); K04263 prostacyclin receptor |
| | Chipl03467 | AKT2; AKT serine/threonine kinase 2; K04456 RAC serine/threonine-protein kinase [EC:2.7.11.1] |
| chr41 | Chipl03468 | TGFB1; transforming growth factor beta 1; K13375 transforming growth factor beta-1 |

| | | |
|---|---|---|
| | Chipl03496 | RASGRP4; RAS guanyl releasing protein 4; K04350 RAS guanyl-releasing protein 1 |
| | Chipl03535 | T-cell receptor beta-1 chain C region; K10785 T-cell receptor beta chain V region |
| | Chipl03536 | T-cell receptor beta-2 chain C region; K10785 T-cell receptor beta chain V region |
| | Chipl03537 | uncharacterized LOC102353650; K10785 T-cell receptor beta chain V region |
| | Chipl03539 | Ig heavy chain Mem5-like; K10785 T-cell receptor beta chain V region |
| | Chipl03541 | C1S; complement C1s; K01331 complement component 1, s subcomponent [EC:3.4.21.42] |
| | Chipl03544 | C1S; complement C1s; K01331 complement component 1, s subcomponent [EC:3.4.21.42] |
| | Chipl03795 | cdk4; cyclin dependent kinase 4; K02089 cyclin-dependent kinase 4 [EC:2.7.11.22] |
| | Chipl03871 | tank; TRAF family member associated NFKB activator; K12650 TRAF family member-associated NF-kappa-B activator |
| chr43 | Chipl03896 | tumor necrosis factor receptor superfamily member 5-like; K03160 tumor necrosis factor receptor superfamily member 5 |
| | Chipl17278 | synaptogenesis protein syg-2-like; K06856 immunoglobulin heavy chain |
| | Chipl17279 | uncharacterized LOC101027262; K06856 immunoglobulin heavy chain |
| | Chipl17395 | uncharacterized LOC102279674; K10784 T cell receptor alpha chain V region |
| | Chipl17397 | uncharacterized LOC102234852; K10784 T cell receptor alpha chain V region |
| chr44 | Chipl17398 | uncharacterized LOC100621222; K10784 T cell receptor alpha chain V region |

| | |
|---|---|
| Chipl17401 | TRAV1J2C, TRAV2J13C, TRAV3J3C; uncharacterized LOC101074543; K10784 T cell receptor alpha chain V region |
| Chipl17403 | T-cell receptor alpha chain V region RL-5; K10784 T cell receptor alpha chain V region |
| Chipl17405 | uncharacterized LOC105234922; K10784 T cell receptor alpha chain V region |
| Chipl17414 | uncharacterized LOC102234852; K10784 T cell receptor alpha chain V region |
| Chipl17415 | uncharacterized LOC101471379; K10784 T cell receptor alpha chain V region |
| Chipl17419 | TRAV14S2; mCG140239-like; K10784 T cell receptor alpha chain V region |
| Chipl17423 | T-cell receptor alpha chain V region CTL-F3; K10784 T cell receptor alpha chain V region |
| Chipl17424 | T-cell receptor alpha chain V region CTL-F3; K10784 T cell receptor alpha chain V region |
| Chipl17425 | uncharacterized LOC710149; K10784 T cell receptor alpha chain V region |
| Chipl17429 | TRAV1J2C, TRAV2J13C, TRAV3J3C; uncharacterized LOC101074543; K10784 T cell receptor alpha chain V region |
| Chipl17430 | tra.L, tcra, tcrd, tra, tra@; T-cell receptor alpha locus L homeolog; K10784 T cell receptor alpha chain V region |
| Chipl17431 | uncharacterized LOC106731799; K06856 immunoglobulin heavy chain |
| Chipl17432 | TRAV1J2C, TRAV2J13C, TRAV3J3C; uncharacterized LOC101074543; K10784 T cell receptor alpha chain V region |
| Chipl17433 | uncharacterized LOC106731799; K06856 immunoglobulin heavy chain |
| Chipl17434 | TRAV1J2C, TRAV2J13C, TRAV3J3C; uncharacterized LOC101074543; K10784 T cell receptor alpha chain V region |
| Chipl17435 | uncharacterized LOC102234852; K10784 T cell receptor alpha chain V region |

| | | |
|---|---|---|
| | Chipl17436 | TRAV1J2C, TRAV2J13C, TRAV3J3C; uncharacterized LOC101074543; K10784 T cell receptor alpha chain V region |
| | Chipl17437 | uncharacterized LOC102149536; K10784 T cell receptor alpha chain V region |
| | Chipl17479 | CFL2; cofilin 2; K05765 cofilin |
| | Chipl17520 | nuclear transcription factor Y subunit C-2-like; K08066 nuclear transcription factor Y, gamma |
| chr45 | Chipl17572 | nfkbie; NFKB inhibitor epsilon; K05872 NF-kappa-B inhibitor epsilon |
| | Chipl12427 | apoptosis-associated speck-like protein containing a CARD; K12799 apoptosis-associated speck-like protein containing a CARD |
| | Chipl12431 | uncharacterized LOC101922302; K10785 T-cell receptor beta chain V region |
| | Chipl12432 | T-cell receptor beta-2 chain C region; K10785 T-cell receptor beta chain V region |
| chr46 | Chipl12448 | CARD6; caspase recruitment domain family member 6; K12797 caspase recruitment domain-containing protein 6 |
| | Chipl17218 | leukocyte immunoglobulin-like receptor subfamily A member 6; K06512 leukocyte immunoglobulin-like receptor |
| | Chipl17232 | urokinase plasminogen activator surface receptor-like; K03985 plasminogen activator, urokinase receptor |
| | Chipl17240 | ICOSLG; inducible T-cell costimulator ligand; K06710 inducible T-cell co-stimulator ligand |
| | Chipl17260 | C-X-C chemokine receptor type 3-like; K04188 C-X-C chemokine receptor type 3 |
| | Chipl17261 | C-X-C chemokine receptor type 3-like; K04188 C-X-C chemokine receptor type 3 |
| chr47 | Chipl17267 | PLAUR; plasminogen activator, urokinase receptor; K03985 plasminogen activator, urokinase receptor |
| chr48 | Chipl03948 | thyroid receptor-interacting protein 6-like; K12792 thyroid receptor-interacting protein 6 |

| | | |
|---|---|---|
| | Chipl03964 | APRIL; K05475 tumor necrosis factor ligand superfamily member 13 |
| | Chipl03973 | CARD6; caspase recruitment domain family member 6; K12797 caspase recruitment domain-containing protein 6 |
| | Chipl03974 | CARD6; caspase recruitment domain family member 6; K12797 caspase recruitment domain-containing protein 6 |
| | Chipl03984 | SERPINE1; serpin family E member 1; K03982 plasminogen activator inhibitor 1 |
| | Chipl03994 | NAIP; NLR family apoptosis inhibitory protein; K12807 baculoviral IAP repeat-containing protein 1 |
| | Chipl03995 | complement C1s subcomponent-like; K01331 complement component 1, s subcomponent [EC:3.4.21.42] |
| | Chipl03996 | mannan-binding lectin serine protease 2-like; K01331 complement component 1, s subcomponent [EC:3.4.21.42] |
| | Chipl04000 | ARRB2; arrestin beta 2; K04439 beta-arrestin |
| | Chipl04011 | epo; erythropoietin; K05437 erythropoietin |
| | Chipl04017 | slit homolog 3 protein-like; K06261 platelet glycoprotein Ib alpha chain |
| | Chipl04019 | GNB2; G protein subunit beta 2; K04537 guanine nucleotide-binding protein G(I)/G(S)/G(T) subunit beta-2 |
| | Chipl04031 | TFRC; transferrin receptor; K06503 transferrin receptor |
| | Chipl04032 | phosphatidylinositol 4,5-bisphosphate 3-kinase catalytic subunit alpha isoform-like; K00922 phosphatidylinositol-4,5-bisphosphate 3-kinase catalytic subunit alpha/beta/delta [EC:2.7.1.153] |
| | Chipl04040 | CLDN15; claudin 15; K06087 claudin |
| | Chipl04056 | pld2; phospholipase D2; K01115 phospholipase D1/2 [EC:3.1.4.4] |
| chr49 | Chipl17447 | PRF1; perforin 1; K07818 perforin 1 |
| chr50 | Chipl02514 | DNA-directed RNA polymerase III subunit RPC2-like; K03021 DNA-directed RNA polymerase III subunit RPC2 [EC:2.7.7.6] |

| | | |
|---|---|---|
| | Chipl02532 | FCGR3A; low affinity immunoglobulin gamma Fc region receptor III; K06463 low affinity immunoglobulin gamma Fc receptor III |
| | Chipl02536 | FCGR3A; low affinity immunoglobulin gamma Fc region receptor III; K06463 low affinity immunoglobulin gamma Fc receptor III |
| | Chipl02538 | FCER1A; Fc fragment of IgE receptor Ia; K08089 Fc receptor, IgE, high affinity I, alpha polypeptide |
| | Chipl02552 | leukocyte immunoglobulin-like receptor subfamily B member 2; K06512 leukocyte immunoglobulin-like receptor |
| | Chipl03255 | PIP5K1A; phosphatidylinositol-4-phosphate 5-kinase type 1 alpha; K00889 1-phosphatidylinositol-4-phosphate 5-kinase [EC:2.7.1.68] |
| | Chipl03264 | GB10346; partitioning defective 3 homolog; K04237 partitioning defective protein 3 |
| | Chipl03271 | interferon gamma receptor 1-like; K05132 interferon gamma receptor 1 |
| chr51 | Chipl03286 | ADAR; adenosine deaminase, RNA specific; K12968 double-stranded RNA-specific adenosine deaminase [EC:3.5.4.37] |

**Supplementary Table 11.** List of MHC class I genes in analyzed species.

| Species | Gene ID | Description from KEGG |
|---|---|---|
| | ENSORLT00000025274 | class I histocompatibility antigen, F10 alpha chain-like; |
| | ENSORLT00000025098 | orla-uga; major histocompatibility complex class I-related gene protein-like; |
| | ENSORLT00000008080 | major histocompatibility complex class I-related gene protein-like; |
| | ENSORLT00000021463 | orla-uha; major histocompatibility complex class I-related gene protein-like; |
| Medaka | ENSORLT00000008540 | orla-uba; major histocompatibility complex class I-related gene protein-like; |
| | ENSORLT00000008514 | orla-uaa; major histocompatibility complex class I-related gene protein-like; |
| | ENSORLT00000023403 | major histocompatibility complex class I-related gene protein-like; |
| | ENSORLT00000025228 | orla-uia1; major histocompatibility complex class I-related gene protein-like; |
| | ENSORLT00000001321 | H-2 class I histocompatibility antigen, L-D alpha chain-like; |
| | ENSORLT00000015537 | UCA; class I histocompatibility antigen, F10 alpha chain-like; |

| | | |
|---|---|---|
| | ENSORLT00000001306 | H-2 class I histocompatibility antigen, L-D alpha chain-like; |
| | ENSORLT00000001185 | major histocompatibility complex class I-related gene protein-like; |
| | ENSORLT00000024360 | major histocompatibility complex class I-related gene protein-like; |
| | ENSORLT00000008255 | Orla-UDA; class I histocompatibility antigen, F10 alpha chain; |
| | ENSORLT00000001202 | major histocompatibility complex class I-related gene protein-like; |
| | ENSORLT00000001304 | class I histocompatibility antigen, F10 alpha chain-like; |
| | ENSLACT00000004968 | Lach-UC-01; major histocompatibility complex class I-related gene protein-like; |
| | ENSLACT00000006461 | Lach-UB-03, Lach-UX-01; uncharacterized LOC102345632; |
| | ENSLACT00000005288 | class I histocompatibility antigen, F10 alpha chain-like; |
| | ENSLACT00000008428 | Lach-UB-05, Lach-UB-06; class I histocompatibility antigen, F10 alpha chain; |
| | ENSLACT00000008716 | Lach-UB-05, Lach-UB-06; class I histocompatibility antigen, F10 alpha chain; |
| | ENSLACT00000013000 | Lach-UA-02; uncharacterized LOC102360671; |
| | ENSLACT00000002454 | Lach-UD-01; major histocompatibility complex class I-related gene protein-like; |
| | ENSLACT00000000716 | major histocompatibility complex class I-related gene protein-like; |
| | ENSLACT00000014038 | class I histocompatibility antigen, F10 alpha chain; |
| | ENSLACT00000006249 | Lach-UA-01; class I histocompatibility antigen, F10 alpha chain-like; |
| | ENSLACT00000015427 | Lach-UA-01; class I histocompatibility antigen, F10 alpha chain-like; |
| | ENSLACT00000000145 | Lach-UD-01; major histocompatibility complex class I-related gene protein-like; |
| Ccoelacanth | ENSLACT00000007335 | Lach-UB-03, Lach-UX-01; uncharacterized LOC102345632; |
| | ENSLACT00000006939 | major histocompatibility complex class I-related gene protein-like; |
| | ENSLACT00000001389 | major histocompatibility complex class I-related gene protein-like; |
| | ENSLACT00000010623 | major histocompatibility complex class I-related gene protein-like; |
| | ENSLACT00000003977 | major histocompatibility complex class I-related gene protein-like; |
| | ENSLACT00000009038 | Lach-UB-03, Lach-UX-01; uncharacterized LOC102345632; |
| | ENSLACT00000011445 | major histocompatibility complex class I-related gene protein-like; |
| | ENSLACT00000003762 | class I histocompatibility antigen, F10 alpha chain-like; |
| | ENSLACT00000000706 | Lach-UD-01; major histocompatibility complex class I-related gene protein-like; |
| | ENSLACT00000008015 | Lach-UB-05; major histocompatibility complex class I-related gene protein-like; |
| | ENSLACT00000008175 | Lach-UD-01; major histocompatibility complex class I-related gene protein-like; |
| | ENSLACT00000003777 | class I histocompatibility antigen, F10 alpha chain; |
| | ENSLACT00000012588 | Lach-UA-02; uncharacterized LOC102360671; |

| | | |
|---|---|---|
| Three-spined sticklebacks | ENSGACT00000000162 | class I histocompatibility antigen, F10 alpha chain; |
| | ENSGACT00000000184 | class I histocompatibility antigen, F10 alpha chain; |
| | ENSGACT00000000156 | class I histocompatibility antigen, F10 alpha chain; |
| | ENSGACT00000000148 | orla-uaa; Mhc class I A; |
| | ENSGACT00000001679 | class I histocompatibility antigen, F10 alpha chain; |
| | ENSGACT00000000197 | class I histocompatibility antigen, F10 alpha chain; |
| | ENSGACT00000002390 | class I histocompatibility antigen, F10 alpha chain; |
| | ENSGACT00000002575 | class I histocompatibility antigen, F10 alpha chain; |
| | ENSGACT00000000165 | class I histocompatibility antigen, F10 alpha chain; |
| | ENSGACT00000002840 | class I histocompatibility antigen, F10 alpha chain; |
| | ENSGACT00000002527 | class I histocompatibility antigen, F10 alpha chain; |
| | ENSGACT00000002518 | major histocompatibility complex class I-related gene protein-like; |
| | ENSGACT00000002485 | class I histocompatibility antigen, F10 alpha chain; |
| | ENSGACT00000002523 | class I histocompatibility antigen, F10 alpha chain; |
| | ENSGACT00000002499 | class I histocompatibility antigen, F10 alpha chain; |
| | ENSGACT00000002577 | class I histocompatibility antigen, Gogo-OKO alpha chain-like; |
| | ENSGACT00000002570 | class I histocompatibility antigen, F10 alpha chain; |
| | ENSGACT00000002491 | class I histocompatibility antigen, F10 alpha chain; |
| | ENSGACT00000002530 | class I histocompatibility antigen, F10 alpha chain; |
| | ENSGACT00000012867 | class I histocompatibility antigen, F10 alpha chain-like; |
| Zebrafish | ENSDART00000133954 | mhc1uea, Dare-UEA, UEA, fb69e07, wu:fb69e07; major histocompatibility complex class I UEA; |
| | ENSDART00000078213 | major histocompatibility complex class I-related gene protein-like; |
| | ENSDART00000143807 | major histocompatibility complex class I-related gene protein-like; |
| | ENSDART00000158560 | major histocompatibility complex class I-related gene protein-like; |
| | ENSDART00000145890 | major histocompatibility complex class I-related gene protein-like; |
| | ENSDART00000160922 | major histocompatibility complex class I-related gene protein-like; |
| | ENSDART00000073383 | major histocompatibility complex class I-related gene protein-like; |
| | ENSDART00000063529 | major histocompatibility complex class I-related gene protein-like; |
| | ENSDART00000161964 | major histocompatibility complex class I-related gene protein-like; |
| | ENSDART00000053153 | major histocompatibility complex class I-related gene protein-like; |

| | | |
|---|---|---|
| | ENSDART00000156529 | major histocompatibility complex class I-related gene protein-like; |
| | ENSDART00000073382 | major histocompatibility complex class I-related gene protein-like; |
| | ENSDART00000164194 | major histocompatibility complex class I-related gene protein-like; |
| | ENSDART00000016845 | major histocompatibility complex class I-related gene protein-like; |
| | ENSDART00000073381 | major histocompatibility complex class I-related gene protein-like; |
| | ENSDART00000154975 | major histocompatibility complex class I-related gene protein-like; |
| | ENSDART00000101143 | major histocompatibility complex class I-related gene protein-like; |
| | ENSDART00000157325 | major histocompatibility complex class I-related gene protein-like; |
| | ENSDART00000057224 | BOLA class I histocompatibility antigen, alpha chain BL3-7-like; |
| | ENSDART00000045182 | major histocompatibility complex class I-related gene protein-like; |
| | ENSDART00000153472 | major histocompatibility complex class I-related gene protein-like; |
| | ENSDART00000082034 | mhc1uea, Dare-UEA, UEA, fb69e07, wu:fb69e07; major histocompatibility complex class I UEA; |
| | ENSDART00000172490 | major histocompatibility complex class I-related gene protein-like; |
| | ENSDART00000009689 | mhc1uea, Dare-UEA, UEA, fb69e07, wu:fb69e07; major histocompatibility complex class I UEA; |
| Whale shark | XP_020365160.1 | class I histocompatibility antigen, F10 alpha chain; |
| | XP_020366167.1 | class I histocompatibility antigen, F10 alpha chain; |
| | XP_020366168.1 | class I histocompatibility antigen, F10 alpha chain; |
| | XP_020370556.1 | mhc1uxa2, MHC1, PAC-BS1, zgc:65799, zgc:77509; major histocompatibility complex class I UXA2 gene; |
| | XP_020370558.1 | Lach-UA-01; class I histocompatibility antigen, F10 alpha chain-like; |
| | XP_020371626.1 | Lach-UA-01; class I histocompatibility antigen, F10 alpha chain-like; |
| | XP_020375020.1 | Lach-UA-02; uncharacterized LOC102360671; |
| | XP_020375131.1 | Lach-UA-02; uncharacterized LOC102360671; |
| | XP_020375404.1 | class I histocompatibility antigen, F10 alpha chain-like; |
| | XP_020377628.1 | Lach-UA-02; uncharacterized LOC102360671; |
| | XP_020379478.1 | orla-uaa; major histocompatibility complex class I-related gene protein-like; |
| | XP_020380267.1 | class I histocompatibility antigen, F10 alpha chain-like; |
| | XP_020381308.1 | Lach-UA-02; uncharacterized LOC102360671; |
| | XP_020381483.1 | Sasa-UBA, UBA; class I histocompatibility antigen, F10 alpha chain-like; |

| | | |
|---|---|---|
| | XP_020384586.1 | Lach-UA-02; uncharacterized LOC102360671; |
| | XP_020384588.1 | BOLA class I histocompatibility antigen, alpha chain BL3-7-like; |
| | XP_020384589.1 | Lach-UA-01; class I histocompatibility antigen, F10 alpha chain-like; |
| | XP_020391952.1 | uncharacterized LOC398628; |
| Bamboo shark | Chipl12832 | Lach-UA-02; uncharacterized LOC102360671; |
| | Chipl12833 | Sasa-UBA, UBA; class I histocompatibility antigen, F10 alpha chain-like; |
| | Chipl12873 | Lach-UA-01; class I histocompatibility antigen, F10 alpha chain-like; |
| | Chipl12895 | class I histocompatibility antigen, F10 alpha chain; |
| | Chipl12897 | class I histocompatibility antigen, F10 alpha chain; |
| | Chipl12898 | Lach-UA-01; class I histocompatibility antigen, F10 alpha chain-like; |
| | Chipl12906 | Lach-UA-02; uncharacterized LOC102360671; |
| | Chipl12912 | Lach-UA-01; class I histocompatibility antigen, F10 alpha chain-like; |
| | Chipl12931 | class I histocompatibility antigen, F10 alpha chain; |
| | Chipl12940 | class I histocompatibility antigen, F10 alpha chain; |
| | Chipl12942 | class I histocompatibility antigen, F10 alpha chain; |
| | Chipl18585 | MODO-UE; MHC class I antigen; |
| | Chipl02590 | class I histocompatibility antigen, F10 alpha chain-like; |
| | Chipl03555 | class I histocompatibility antigen, F10 alpha chain; |
| | Chipl14141 | class I histocompatibility antigen, F10 alpha chain-like; |
| | Chipl18471 | class I histocompatibility antigen, F10 alpha chain; |
| | Chipl19068 | Lach-UA-02; uncharacterized LOC102360671; |
| | Chipl19417 | Lach-UA-01; class I histocompatibility antigen, F10 alpha chain-like; |
| Elephant shark | LOC103189724 | major histocompatibility complex class I-related gene protein-like |
| | LOC103173525 | class I histocompatibility antigen, F10 alpha chain-like |
| | LOC103174402 | major histocompatibility complex class I-related gene protein-like |
| | LOC103173953 | class I histocompatibility antigen, F10 alpha chain-like |
| | LOC103173736 | class I histocompatibility antigen, F10 alpha chain-like |

**Supplementary Table 12.** List of MHC class II genes in analyzed species.

| Species | Gene ID | Description from KEGG |
|---|---|---|
| Medaka | ENSORLT00000000027 | HLA class II histocompatibility antigen, DP alpha 1 chain-like; |

| | | |
|---|---|---|
| | ENSORLT00000016063 | rano class II histocompatibility antigen, A beta chain-like; |
| | ENSORLT00000005537 | CD74 molecule, major histocompatibility complex, class II invariant chain a |
| | ENSORLT00000016021 | Orla-DDA; RLA class II histocompatibility antigen, DP alpha-1 chain; |
| | ENSORLT00000000030 | Orla-DCB; H-2 class II histocompatibility antigen, E-S beta chain; |
| | ENSORLT00000023575 | Orla-DAA; mamu class II histocompatibility antigen, DR alpha chain; |
| | ENSORLT00000023543 | Orla-DAB; H-2 class II histocompatibility antigen, E-S beta chain; |
| | ENSORLT00000011500 | Orla-DEA; RLA class II histocompatibility antigen, DP alpha-1 chain-like; |
| | ENSORLT00000016052 | Orla-DDB; rano class II histocompatibility antigen, A beta chain; |
| | ENSORLT00000024164 | Orla-DFA; mamu class II histocompatibility antigen, DR alpha chain; |
| | ENSORLT00000006056 | H-2 class II histocompatibility antigen, A-U alpha chain-like; |
| | ENSORLT00000024129 | H-2 class II histocompatibility antigen, E-S beta chain-like; |
| Coelacanth | ENSLACT00000001762 | H-2 class II histocompatibility antigen, E-S beta chain-like; |
| | ENSLACT00000000411 | DLA class II histocompatibility antigen, DR-1 beta chain-like; |
| | ENSLACT00000000328 | SLA class II histocompatibility antigen, DQ haplotype C beta chain-like; |
| | ENSLACT00000000754 | HLA class II histocompatibility antigen, DP alpha 1 chain-like; |
| | ENSLACT00000000891 | rano class II histocompatibility antigen, D-1 beta chain-like; |
| | ENSLACT00000003230 | RLA class II histocompatibility antigen, DP alpha-1 chain-like; |
| | ENSLACT00000000197 | H-2 class II histocompatibility antigen, A-D beta chain-like; |
| | ENSLACT00000000493 | DLA class II histocompatibility antigen, DR-1 beta chain-like; |
| | ENSLACT00000000162 | rano class II histocompatibility antigen, D-1 beta chain-like; |
| | ENSLACT00000002416 | RLA class II histocompatibility antigen, DP alpha-1 chain-like; |
| | ENSLACT00000002328 | DLA class II histocompatibility antigen, DR-1 beta chain-like; |
| | ENSLACT00000009069 | DLA class II histocompatibility antigen, DR-1 beta chain-like; |
| | ENSLACT00000004598 | SLA class II histocompatibility antigen, DQ haplotype C beta chain-like; |
| | ENSLACT00000000968 | H-2 class II histocompatibility antigen, A-B alpha chain-like; |
| | ENSLACT00000000721 | H-2 class II histocompatibility antigen, A-U alpha chain-like; |
| | ENSLACT00000000561 | DLA class II histocompatibility antigen, DR-1 beta chain-like; |
| | ENSLACT00000008726 | DLA class II histocompatibility antigen, DR-1 beta chain-like; |
| | ENSLACT00000000289 | SLA class II histocompatibility antigen, DQ haplotype C beta chain-like; |
| | ENSLACT00000006607 | H-2 class II histocompatibility antigen, E-S beta chain-like; |
| | ENSGACT00000000421 | H-2 class II histocompatibility antigen, A-U alpha chain-like; |

| | | |
|---|---|---|
| Three-spined sticklebacks | ENSGACT00000025242 | Orla-DAA; mamu class II histocompatibility antigen, DR alpha chain; |
| | ENSGACT00000000431 | H-2 class II histocompatibility antigen, E-S beta chain-like; |
| | ENSGACT00000004860 | rano class II histocompatibility antigen, A beta chain-like; |
| | ENSGACT00000025238 | Orla-DAB; H-2 class II histocompatibility antigen, E-S beta chain; |
| | ENSGACT00000000450 | Orla-DAB; H-2 class II histocompatibility antigen, E-S beta chain; |
| | ENSGACT00000000437 | HLA class II histocompatibility antigen, DRB1-8 beta chain-like; |
| | ENSGACT00000023783 | H-2 class II histocompatibility antigen, E-S beta chain-like; |
| | ENSGACT00000004910 | H-2 class II histocompatibility antigen, A-U alpha chain-like; |
| | ENSGACT00000000439 | Orla-DAA; mamu class II histocompatibility antigen, DR alpha chain; |
| | ENSGACT00000000434 | H-2 class II histocompatibility antigen, A-U alpha chain-like; |
| Zebrafish | ENSDART00000006898 | novel protein with a Class II histocompatibility antigen, alpha domain and a Immunoglobulin C1-set domain; |
| | ENSDART00000161194 | MHCII, si:busm1-48c11.4, si:dz194e12.12, si:dz48c11.4, zgc:123061; si:busm1-194e12.12; |
| | ENSDART00000102847 | si:dz48c11.3; si:busm1-48c11.3; |
| | ENSDART00000160609 | mhc2dab, major histocompatibility complex class II DAB gene; |
| | ENSDART00000129901 | novel protein with a Class II histocompatibility antigen, alpha domain and a Immunoglobulin C1-set domain; |
| | ENSDART00000109439 | MHCII, si:busm1-160c18.10, si:dz160c18.10, si:dz194e12.8, zgc:101701; si:busm1-194e12.8; |
| | ENSDART00000099281 | mhc2dcb, major histocompatibility complex class II DCB gene; |
| | ENSDART00000097932 | MHCII, si:busm1-48c11.4, si:dz194e12.12, si:dz48c11.4, zgc:123061; si:busm1-194e12.12; |
| | ENSDART00000159361 | MHCII, si:busm1-48c11.4, si:dz194e12.12, si:dz48c11.4, zgc:123061; si:busm1-194e12.12; |
| | ENSDART00000162877 | si:dz48c11.3; si:busm1-48c11.3; |
| | ENSDART00000159276 | MHCII, si:busm1-48c11.4, si:dz194e12.12, si:dz48c11.4, zgc:123061; si:busm1-194e12.12; |
| | ENSDART00000123658 | mhc2dcb, si:busm1-243a08.1, si:busm1-37i06.8, si:dz243a08.1, si:dz37i06.8; major histocompatibility complex class II DCB gene; |
| | ENSDART00000053205 | CD74 molecule, major histocompatibility complex, class II invariant chain a; |

| | | |
|---|---|---|
| | ENSDART00000168831 | mhc2dab, major histocompatibility complex class II DAB gene; |
| | ENSDART00000168076 | mhc2dab,major histocompatibility complex class II DAB gene; |
| | ENSDART00000171760 | si:dz48c11.3; si:busm1-48c11.3; |
| | ENSDART00000026021 | CD74 molecule, major histocompatibility complex, class II invariant chain a; |
| | ENSDART00000158400 | MHCII, si:busm1-160c18.10, si:dz160c18.10, si:dz194e12.8, zgc:101701; si:busm1-194e12.8; |
| | ENSDART00000158784 | mhc2dcb, si:busm1-243a08.1, si:busm1-37i06.8, si:dz243a08.1, si:dz37i06.8; major histocompatibility complex class II DCB gene; |
| | ENSDART00000048448 | uncharacterized LOC791723; |
| | ENSDART00000105025 | H-2 class II histocompatibility antigen, E-D beta chain-like; |
| Whale shark | XP_020390554.1 | ORAN-DRA; MHC class II DR alpha; |
| | XP_020390555.1 | DRB, DRB1, Mamu-DRB, Mamu-DRB1, Mane-DRB; HLA class II histocompatibility antigen, DRB1-3 chain; |
| | XP_020392206.1 | DLA class II histocompatibility antigen, DR-1 beta chain-like; |
| Bamboo shark | Chipl12839 | DLA class II histocompatibility antigen, DR-1 beta chain-like; |
| | Chipl12840 | RLA class II histocompatibility antigen, DP alpha-1 chain-like; |
| | Chipl12907 | RLA class II histocompatibility antigen, DP alpha-1 chain-like; |
| | Chipl01708 | class II histocompatibility antigen, B-L beta chain-like; |
| | Chipl14140 | SAHA-DAB; HLA class II histocompatibility antigen, DRB1-15 beta chain; |
| | Chipl18694 | HLA class II histocompatibility antigen, DR alpha chain-like; |

**Supplementary Table 13.** MHC class II fragments detected by using BLAST in the elephant shark genome**.**

| Target scaffolds | Query sequences | Identity | Target Start | Target End | Query start | Query End | E value |
|---|---|---|---|---|---|---|---|
| NW_006895284.1 | ENSGACT00000025242 | 93.55 | 2641 | 2671 | 32 | 2 | 2.00E-06 |
| NW_006895810.1 | ENSGACT00000025242 | 93.55 | 3175 | 3205 | 32 | 2 | 2.00E-06 |
| NW_006896611.1 | ENSGACT00000025242 | 93.55 | 2160 | 2190 | 2 | 32 | 2.00E-06 |
| NW_006896785.1 | ENSGACT00000025242 | 93.55 | 283 | 313 | 2 | 32 | 2.00E-06 |
| NW_006896886.1 | ENSGACT00000025242 | 93.55 | 1221 | 1251 | 32 | 2 | 2.00E-06 |
| NW_006898245.1 | Chipl12840 | 86.96 | 355 | 400 | 299 | 344 | 4.00E-06 |
| NW_006898335.1 | ENSGACT00000025242 | 93.33 | 175 | 204 | 32 | 3 | 4.00E-06 |
| NW_006900613.1 | ENSGACT00000025242 | 93.55 | 354 | 384 | 2 | 32 | 9.00E-07 |

| | | | | | | | |
|---|---|---|---|---|---|---|---|
| NW_006910024.1 | ENSGACT00000025242 | 93.55 | 622 | 652 | 32 | 2 | 6.00E-07 |

**Supplementary Table 14.** Ancestral P2X gene found in amphioxus and ascidiacea genomes.

| Name/Gene ID | Description |
|---|---|
| LOC109481458 | P2X purinoceptor 7-like [Branchiostoma belcheri (Belcher's lancelet)] |
| LOC109474372 | P2X purinoceptor 7-like [Branchiostoma belcheri (Belcher's lancelet)] |
| LOC109472980 | P2X purinoceptor 7-like [Branchiostoma belcheri (Belcher's lancelet)] |
| LOC109464342 | P2X purinoceptor 4-like [Branchiostoma belcheri (Belcher's lancelet)] |
| LOC109462847 | P2X purinoceptor 7-like [Branchiostoma belcheri (Belcher's lancelet)] |
| LOC109462011 | P2X purinoceptor 7-like [Branchiostoma belcheri (Belcher's lancelet)] |
| BRAFLDRAFT_84310(P2RX4) | hypothetical protein [Branchiostoma floridae (Florida lancelet)] |
| BRAFLDRAFT_103732 | hypothetical protein [Branchiostoma floridae (Florida lancelet)] |
| LOC101242976 | P2X purinoceptor 7-like [Ciona intestinalis (vase tunicate)] |
| LOC101242780 | P2X purinoceptor 7-like [Ciona intestinalis (vase tunicate)] |

**Supplementary Table 15.** Protein length of P2X genes in bony fishes and sea lamprey.

| | **Gene ID** | **Protein Length(aa)** | **Gene Name** |
|---|---|---|---|
| | ENSLACT00000004839 | 399 | P2RX3 |
| | ENSLACT00000021101 | 403 | P2RX1 |
| | ENSLACT00000019925 | 388 | P2RX2 |
| Coelacanth | ENSLACT00000025971 | 134 | P2RX7 |
| | ENSLACT00000001552 | 363 | P2RX5 |
| | ENSLACT00000002031 | 419 | P2RX6 |
| | ENSLACT00000024921 | 391 | P2RX4 |
| | ENSDART00000167907 | 597 | p2rx7 |
| | ENSDART00000002866 | 398 | p2rx1 |
| | ENSDART00000098969 | 401 | p2rx4b |
| | ENSDART00000019461 | 400 | p2rx2 |
| Zebrafish | ENSDART00000130628 | 398 | p2rx4a |
| | ENSDART00000171919 | 482 | p2rx5 |
| | ENSDART00000163914 | 411 | p2rx3a |
| | ENSDART00000133926 | 412 | p2rx3b |
| | ENSDART00000029063 | 391 | p2rx5 |

| | | | |
|---|---|---|---|
| Medaka | ENSORLT00000014760 | 397 | p2rx1 |
| | ENSORLT00000004381 | 405 | p2rx3 |
| | ENSORLT00000019713 | 392 | p2rx4 |
| | ENSORLT00000015083 | 458 | p2rx5 |
| | ENSORLT00000009830 | 412 | p2rx2 |
| Sea lamprey | PMZ_0041355-RA_PMZ | 79 | p2rx5.L |
| | PMZ_0041356-RA_PMZ | 205 | P2RX5 |
| | PMZ_0048630-RA_PMZ | 169 | p2rx5.S |
| | PMZ_0048646-RA_PMZ | 160 | p2rx5.S |
| | PMZ_0037740-RA_PMZ | 225 | P2RX5 |
| | PMZ_0034936-RA_PMZ | 160 | p2rx5.S |
| | PMZ_0048622-RA_PMZ | 160 | p2rx5.S |
| | PMZ_0046842-RA_PMZ | 74 | p2rx5 |
| | PMZ_0036668-RA_PMZ | 99 | p2rx5.S |
| | PMZ_0035061-RA_PMZ | 246 | P2RX5 |
| | PMZ_0048786-RA_PMZ | 97 | p2rx5.S |
| | PMZ_0037023-RA_PMZ | 98 | p2rx5.S |
| | PMZ_0036851-RA_PMZ | 244 | P2RX5 |
| | PMZ_0048482-RA_PMZ | 154 | P2RX5 |

**Supplementary Table 16.** Target sites for P2X genes knockout experiments.

* Bold letters indicate the protospacer adjacent motif for CRISPR/Cas9 system.

| Genes for KO | Target sites | |
|---|---|---|
| P2RX3 | T1 | GTCTGTTGGAATAATCAACC**GGG**[*] |
| P2RX5 | T1 | CAGGAGCATGGCTCAGACCT**GGG** |
| P2RX5 | T2 | CGGTGATCGGATATCTGATC**GGG** |

**Supplementary Table 17.** Primers used in P2X genes knockout experiments.

| Primers | Sequences |
|---|---|
| P2RX3F | AGTGAGGAGCGTGTTGGTTC |
| P2RX3R | GGATGGACTTCAGTATAGGGTCAAG |
| P2RX5F | ACCGCGTAATTTCTCCCTGT |

| | |
|---|---|
| P2RX5R | ACAGCACGTCATTCAGGTCA |
| P2RX5F | ACCGCGTAATTTCTCCCTGT |
| P2RX5R | ACAGCACGTCATTCAGGTCA |

1. J. Salse, M. Abrouk, F. Murat, U. M. Quraishi, C. Feuillet, Improved criteria and comparative genomics tool provide new insights into grass paleogenomics. *Briefings in bioinformatics* **10**, 619-630 (2009).
